# Microenvironmental correlates of immune checkpoint inhibitor response in human melanoma brain metastases revealed by T cell receptor and single-cell RNA sequencing

**DOI:** 10.1101/2021.08.25.456956

**Authors:** Christopher A. Alvarez-Breckenridge, Samuel C. Markson, Jackson H. Stocking, Naema Nayyar, Matthew Lastrapes, Matthew R. Strickland, Albert E. Kim, Magali de Sauvage, Ashish Dahal, Juliana M Larson, Joana L. Mora, Andrew W. Navia, Benjamin M. Kuter, Corey M. Gill, Mia Solana Bertalan, Brian Shaw, Alexander Kaplan, Megha Subramanian, Aarushi Jain, Swaminathan Kumar, Husain Danish, Michael White, Osmaan Shahid, Kristen E. Pauken, Brian C. Miller, Dennie T. Frederick, Christine Herbert, McKenzie Shaw, Maria Martinez-Lage, Matthew P. Frosch, Nancy Wang, Elizabeth R. Gerstner, Brian V. Nahed, William T. Curry, Bob S. Carter, Daniel P. Cahill, Genevieve Marie Boland, Benjamin Izar, Michael Davies, Arlene Sharpe, Mario L. Suvà, Ryan J. Sullivan, Priscilla K. Brastianos, Scott L. Carter

**Affiliations:** Departments of Neurosurgery, The University of Texas MD Anderson Cancer Center, Houston, TX, USA; Department of Neurosurgery, Massachusetts General Hospital, Boston, MA, USA; Department of Immunology, Blavatnik Institute, Harvard Medical School, Boston, MA, USA; Evergrande Center for Immunological Diseases, Harvard Medical School and Brigham and Women’s Hospital, Boston, MA, USA; Broad Institute, Harvard University & Massachusetts Institute of Technology, Cambridge, MA, USA; Department of Data Sciences, Dana-Farber Cancer Institute, Boston, MA, USA; Department of Medicine, Harvard Medical School & Massachusetts General Hospital, Boston, MA, USA; Department of Epidemiology, The University of Texas MD Anderson Cancer Center, Houston, TX, USA; Massachusetts General Hospital Cancer Center, Boston, MA, USA; Department of Chemistry, Massachusetts Institute of Technology, Cambridge, MA, USA; Institute for Medical Engineering & Science, Massachusetts Institute of Technology, Cambridge, MA, USA; Koch Institute for Integrative Cancer Research, Massachusetts Institute of Technology, Cambridge, MA, USA; Ragon Institute, Harvard University, Massachusetts Institute of Technology, & Massachusetts General Hospital, Cambridge, MA, USA; Department of Melanoma Medical Oncology, The University of Texas MD Anderson Cancer Center, Houston, TX, USA; Department of Neurology, Memorial Sloan Kettering Cancer Center, New York, NY, USA; Weill Cornell Medical Center, New York, New York, USA; Department of Medical Oncology, Dana-Farber Cancer Institute, Boston, MA, USA; Division of Surgical Oncology, Massachusetts General Hospital, Harvard Medical School, Boston, MA, USA; Department of Pathology and Center for Cancer Research, Massachusetts General Hospital and Harvard Medical School, Boston, MA, USA; C. S. Kubik Laboratory for Neuropathology, Mass General Hospital and Harvard Medical School, Boston, MA, USA; Division of Hematology and Oncology, Columbia University Irving Medical Center, New York, NY, USA; Columbia Center for Translational Immunology, New York, NY, USA; Division of Computational Biology, Dana-Farber Cancer Institute, Boston, MA, USA

## Abstract

Melanoma-derived brain metastases (MBM) represent an unmet clinical need due to central nervous system (CNS) progression as a frequent, end-stage site of disease. Immune checkpoint inhibition (ICI) represents a clinical opportunity against MBM; however, the MBM tumor microenvironment (TME) has not been fully elucidated in the context of ICI. To dissect unique MBM-TME elements and correlates of MBM-ICI response, we collected 32 fresh MBM and performed single cell RNA sequencing of the MBM-TME and T cell receptor clonotyping on T cells from MBM and matched blood and extracranial lesions. We observed myeloid phenotypic heterogeneity, most notably multiple distinct neutrophil states including an IL-8 expressing population that correlated with malignant cell epithelial-to-mesenchymal transition. Additionally, we observe significant relationships between intracranial T cell phenotypes and the distribution of T cell clonotypes intracranially and peripherally. We found that the phenotype, clonotype, and overall number of MBM-infiltrating T cells were associated with response to ICI, suggesting that ICI-responsive MBMs interact with peripheral blood in a manner similar to extracranial lesions. These data demonstrate unique features of the MBM-TME, which may represent potential targets to improve clinical outcomes for patients with MBM.

## Introduction

Brain metastases, arising most commonly from lung, breast and melanoma^1, 2^, represent the most common type of intracranial tumor, occurring in 20-40% of patients diagnosed with cancer^2–8^. Intracranial progression frequently occurs even within the context of extracranial response to therapy and nearly half of patients with symptomatic brain metastases succumb to their disease ^9^. Metastatic tumors disseminating into the central nervous system (CNS) are associated with a poor prognosis and have traditionally been relegated to surgical resection and radiotherapy. Within this standard of care, intracranial metastases have been associated with significant morbidity and median survival ranges from 3 to 27 months^10^. In the setting of their increasing prevalence, limited treatment options, and historical exclusion from clinical trials, brain metastases represent an unmet clinical need.

Immune checkpoint inhibitors (ICI) have revolutionized the treatment of cancer with approval in 19 cancer types and two tissue-agnostic indications^11^. With the growing clinical success of immune checkpoint modulation, attention has shifted towards the potential activity of ICI in treating brain metastases. Studies targeting CTLA-4 and PD-1 have demonstrated promising ICI mediated intracranial response rates for metastatic melanoma, with combination therapy achieving similar rates of response observed extracranially^12–15^. Beyond melanoma brain metastases, intracranial efficacy has similarly been demonstrated with pembrolizumab in patients with renal cell carcinoma^16^ and non-small-cell lung cancer (NSCLC)^13^.

Despite these promising clinical results, approximately half of all melanoma patients progress on ICI secondary to either innate or acquired resistance. While extracranial disease progression occurs through both tumor-intrinsic^17^ and extrinsic^18^ mechanisms, a paucity of data exists investigating determinants of intracranial response and/or resistance to ICI. The CNS features an immune specialized microenvironment^19–22^ and the interaction of these CNS-unique elements with ICI is not fully understood. Clinical evidence suggests that the presence of extracranial lesions influences immune based therapies for brain metastases^8, 15, 23^. Potential mechanisms for this process include T cell priming at extracranial sites of disease and T cell trafficking into the brain where shared intra- and extracranial tumor antigens are targeted.

Several features of response and resistance to ICI in the extracranial setting have been investigated. At a genomic level, the overall tumor mutational burden has been associated with clinical response to anti-PD1 therapy^24, 25^ in addition to predicted neoantigen load at pre-treatment timepoints^26^. Additionally, many studies have investigated the role of T cell infiltration prior to ICI^27, 28^, the spatial distribution of CD8+ T cells along tumor margins ^27^, and the extent of PD-L1 expression within the tumor^29, 30^.

However, the results of these findings do not provide a definitive link to ICI response. With the advent of single cell sequencing techniques to dissect immune cell phenotypes, further insights have been made into phenotypic T cell states that are enriched within responding tumors^31^. In order to explore these features within the intracranial context, we utilized single cell RNA sequencing (scRNA-seq) from a cohort of immunotherapy naive and post-ICI treated melanoma-derived brain metastases (MBM) paired with T cell receptor sequencing (TCR-seq) from patient-matched blood and extracranial lesions to describe the diversity of immune and malignant cellular elements within the TME; to identify associations between those elements; and to suggest TME-related biomarkers of intracranial response to ICI.

## RESULTS

### Characterization of the melanoma brain metastatic microenvironment using scRNA-seq

In order to dissect the tumor microenvironment of MBM, we performed scRNA-seq on 32 sequentially collected MBM from 27 unique patients (Fig. 1a,b; Supplementary Table 1). Two patients underwent simultaneous intracranial resections and longitudinal samples were obtained from 2 patients (one contributing 2 samples, the other 3). Of the resected tumors, 23 had been previously treated with ICI, while 9 were immunotherapy naive. Among the ICI-naive individuals, one went on to receive ICI and was a responder both intra- and extracranially. Although the 23 post-ICI patients ultimately experienced intracranial progression resulting in the need for craniotomy, they were defined as having either complete non-response (8) or partial response (15) based on their clinical history of systemic response to ICI. Additionally, 5 patients were treated with targeted therapy prior to resection. Key genetic features of patients’ tumors derived from whole exome sequencing (WES) and the SNaPshot system for SNP genotype are provided in Supplementary Fig. 1.

**Figure 1.**
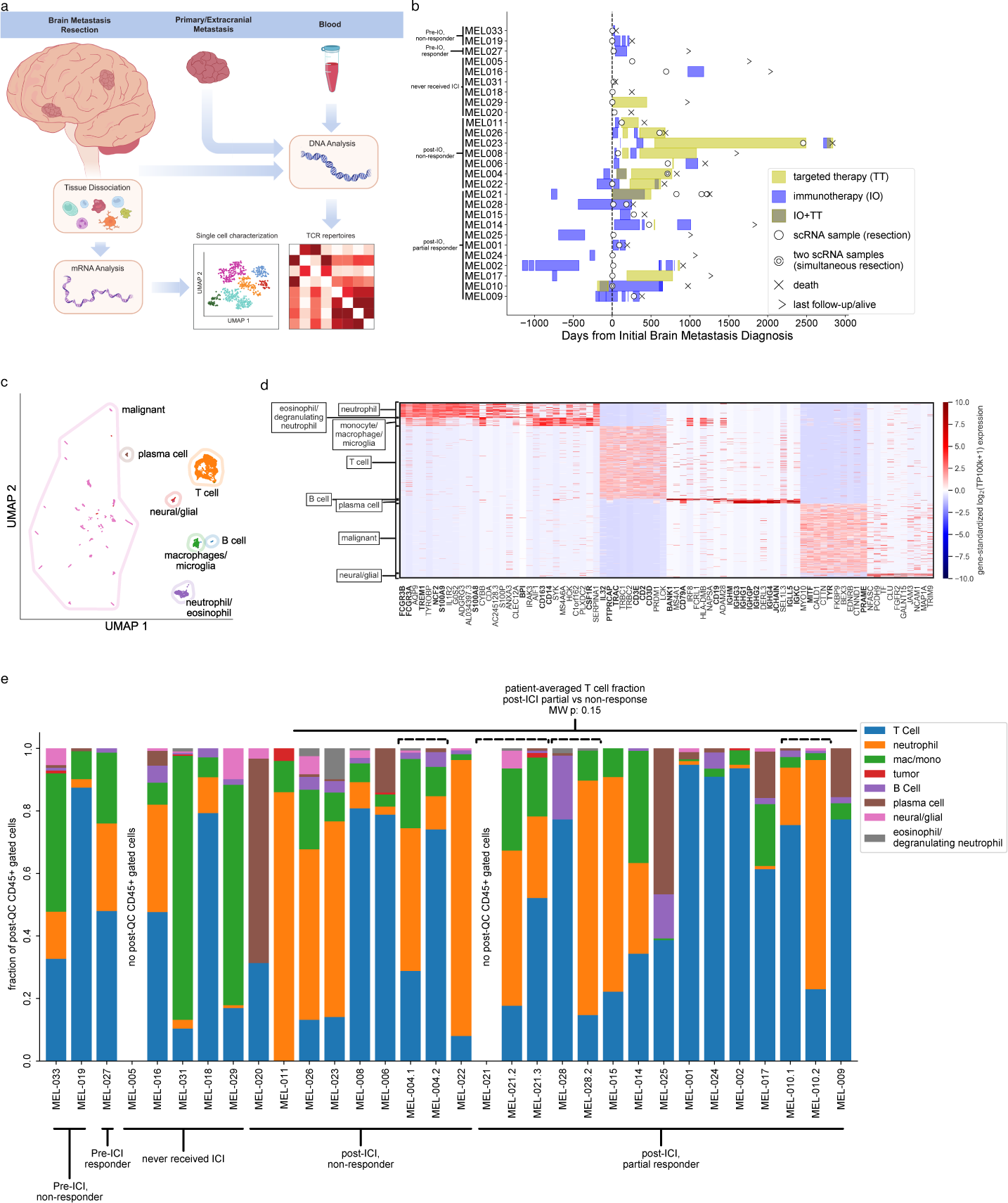
Study design, cohort overview, high-level cell classification and populations. a) Schematic representation of the study. b) Patient clinical trajectories, including timing of immunotherapy and targeted therapy relative to initial diagnosis of brain metastasis. c) UMAP of single-cell transcriptomes, colored and circled by cell type. d) Heatmap of standardized gene expression of key marker genes for each cluster (10 shown per cluster), with notable genes in bold. e) Fraction of post-QC flow cytometry sorted CD45+ cells in each cluster, for each sample. Patient ICI response indicated. Mann-Whitney U p-value is the patient-averaged T cell fraction for post-ICI non-responders vs post-ICI partial responders. Samples derived from the same patient are grouped, with groups indicated by dashed brackets.

Following resection, cells were immediately dissociated and sorted via flow cytometry (see **Methods**) with gating chosen to enrich CD45-, CD45+, and CD45+CD3+ groups. Details regarding dissociation, cDNA generation, and library preparation are described further in **Methods**. In total, we collected and sequenced 19,968 cells using the full transcript length Smart-seq2 protocol ^32^; 14,021 of these cells passed quality control (QC), wherein cells were subsetted based on a minimum unique gene count of 1000 and then clustered via agglomerative clustering, with removal of clusters showing evidence of contamination or doublets (see **Methods**). A median of 2,662, 2,773, and 6,431 unique genes were detected per cell in CD3+, CD45+, and CD45- populations, respectively. Post-QC clusters were also assessed for evidence of cellular stress (Fig. S2)^33^.

Our analysis identified a total of 27 non-immune clusters (see **Methods**). As has previously been reported, malignant cells’ RNA expression was highly patient and sample specific (Fig. 1c)^34–36^. In contrast, the remaining clusters demonstrated cell-type specific gene expression (Fig. 1c,d) profiles including glial cells, lymphoid (T/NK cells, B cells, and plasma cells) and myeloid cells (neutrophil, macrophage/microglia). Relative proportions of CD45+ cells across samples are shown in Fig. 1e. Neither malignant cell PD-L1 expression nor tumor mutational burden were significantly associated with partial vs non-response among post-treatment individuals (Fig. S3).

### Intracranial myeloid populations in the metastatic melanoma TME display unique phenotypes and associations with malignant cell behavior

Utilizing our CD45+ sorted cells, we identified a total of 1,266 neutrophil and 653 monocyte-derived cells (including macrophages and microglia) after QC. Highly expressed genes associated with the macrophage/microglia cluster included *CD14*, *CD163*, and *CSF1R*, while the gene expression profile of neutrophils featured *S100A8*, *S100A9*, and *NCF2* (Fig 1d). Each myeloid population included a cross section of patients from the cohort (Fig. S4a,b).

Within monocyte-derived cells, we observe a cell cluster characterized by elevated relative expression of *S100A8*, *S100A9*, *S100A12*, *MNDA* and a separate cluster of microglia defined by expression of *TREM2*, *APOE*, *C1QA*, *C1QC* (Fig. 2a,b). Canonical microglia markers *TMEM19* and *P2RY12* were additionally upregulated in this cluster (Fig. S5a-d). This latter population of cells has been described elsewhere as reactive microglia, with an elevated presence in the brain parenchyma of individuals with conditions such as multiple sclerosis and Alzheimer’s disease ^37, 38^.

**Figure 2.**
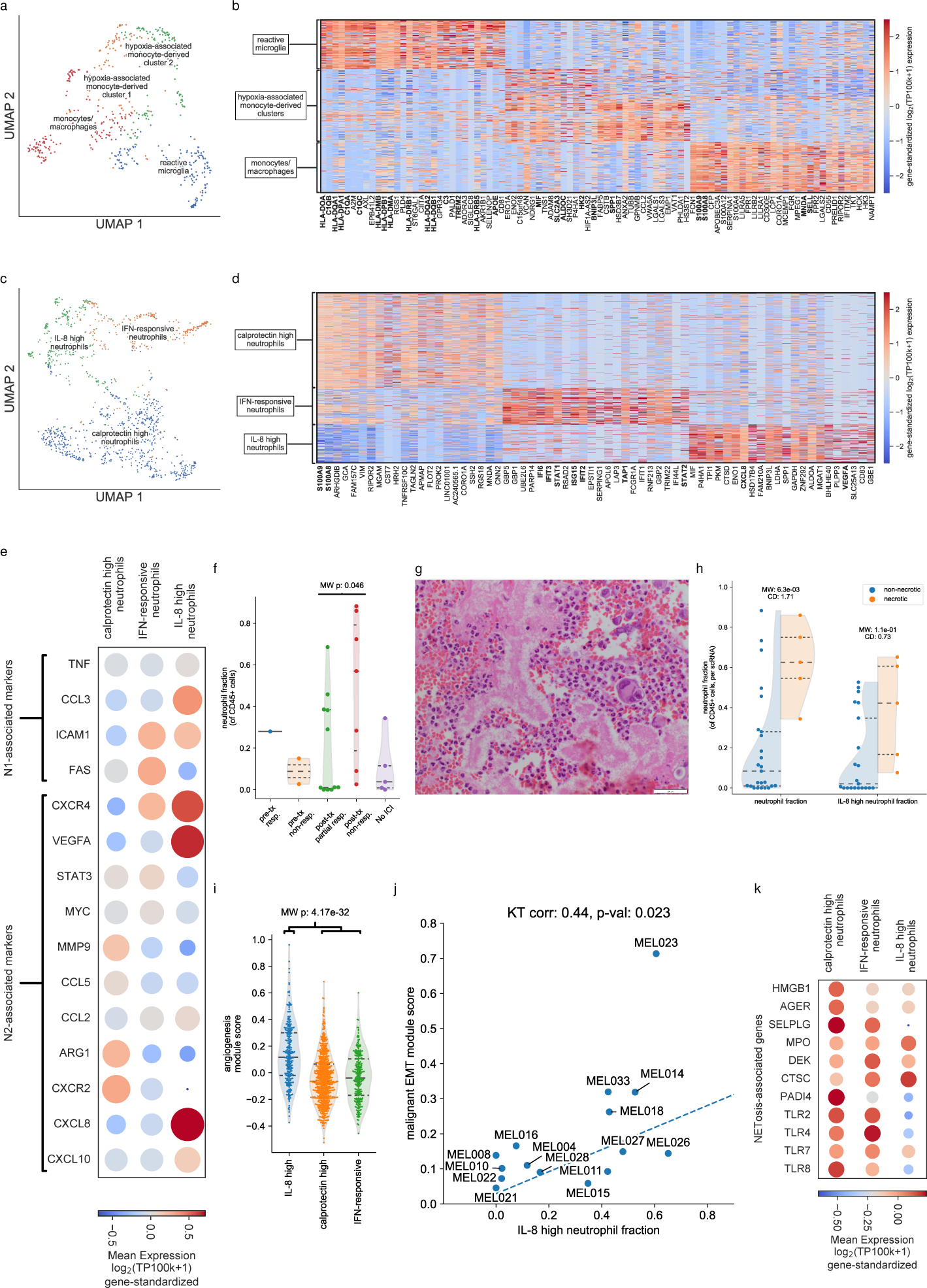
Multiple myeloid phenotypes and their association with malignant phenotypes observed intracranially. a) UMAP of monocyte-derived cells, including macrophages and microglia. b) Heatmap of standardized gene expression of marker genes for clusters of monocyte-derived cells (30 marker genes for reactive microglia and monocytes/macrophages, 15 for each of the hypoxic clusters), with key genes in bold. c) UMAP of neutrophils (excluding degranulating neutrophils/eosinophils). d) Heatmap of standardized gene expression of marker genes for clusters of neutrophils (30 genes per cluster) with key genes in bold. e) Dot plot (see **Methods**) of key genes associated with N1 and N2 phenotypes in the calprotectin high, IFN-responsive, and IL- 8 high neutrophils. f) Distribution (points indicating individual patients are overlaid on kernel density estimate of overall distribution) of fraction of non-degranulating/eosinophil neutrophils (of CD45+ cells) across patients (values for patients contributing multiple samples are averaged); Mann-Whitney U p-value for comparison of post-treatment partial vs non-responders indicated. g) Hematoxylin and eosin (H&E) stain of tumor section from MEL022, showing high levels of blood product and neutrophil infiltration. h) Distribution (points indicating individual patients are overlaid on kernel density estimate of overall distribution) of neutrophil fraction (of post-QC CD45+ cells), IL-8 fraction (of non-degranulating neutrophils) in samples with and without evidence of necrosis. Two-sided Mann-Whitney U p-values indicated. i) Distribution (points indicating individual patients are overlaid on kernel density estimate of overall distribution) of the hallmark angiogenesis module score (see **Methods**) for the IL-8, calprotectin high, and IFN-responsive neutrophils, with Mann-Whitney p-value of IL-8 vs other neutrophils indicated. j) Fraction of IL-8 high neutrophils (of all non- degranulating/eosinophil) vs. EMT module score calculated in malignant cells (see **Methods**) across patients (values for patients contributing multiple samples are averaged); Kendall-τ correlation and associated p-values (see **Methods**) are indicated. Theil-sen line of best fit (see **Methods**) indicated by dotted line. k) Dot plot (see **Methods**) of genes associated with NETosis across calprotectin high, IFN-responsive, and IL-8 high neutrophils.

A total of 1,266 neutrophils were identified in 23 of 27 patients’ cells and were associated with one of four subclusters. One of these subclusters (representing 63 cells from 10 patients) clustered most distantly from the other three neutrophil groups (Fig. S6a). Separate clustering of these cells revealed four clusters (Fig. S6b), with gene expression characteristic of eosinophils (*CCR3*), primary/azurophilic (*MPO*, *AZU1*, *CTSG, DEFA1B*, *DEFA3*, *DEFA4*), secondary/specific (*LTF*, *CAMP*), and tertiary degranulation (*MMP9*), respectively (Fig. S6c,d). These eosinophils/degranulating neutrophils were removed prior to re-clustering the remaining neutrophils. Among the three remaining neutrophil clusters, one included a “calprotectin-high” group characterized by high expression of *S100A8* and *S100A9*, jointly coding for the calprotectin heterodimer, and representing 52% of all identified neutrophils (Fig. 2c,d). A second cluster was termed the “IFN-responsive” neutrophils, which were characterized by high expression of genes associated with interferon gamma response, including *IFI6*, *IFIT2*, *ISG15*, and *TAP1* (Fig. 2c,d). The final cluster was termed the “IL-8-high” neutrophils and was characterized by elevated expression of IL-8/*CXCL8* and *VEGFA* (Fig. 2c,d). The calprotectin-high, IFN-responsive, and IL-8-high neutrophils were observed in 23, 15, and 15 patients, respectively (Fig. S4b).

This neutrophil polarization is partially concordant with what has been seen in murine models and human *in vitro* studies^39, 40^. Previous work on heterogeneity of tumor-associated neutrophils (TANs) has subdivided neutrophils along an “N1” and “N2” polarization axis, corresponding to anti-tumor and pro-tumor phenotypes, respectively^39^. N1 neutrophils were characterized by higher levels of ICAM-1/CD54, FasR/CD95, and TNF- α, whereas N2 neutrophils expressed higher levels of *CCL5*, *CCL2*, *VEGFA*, arginase, and IL-8/*CXCL8*^39, 41^. IL-8-high neutrophils derived from our MBM cohort corresponded most closely to N2 neutrophils; however, they showed reduced expression of N2 markers *MMP9*, and *ARG1*, as well as elevated expression of the N1 markers *CCL3,* and *ICAM1* (Fig. 2e). Therefore, while our data recapitulates the heterogeneous nature of TANs, further studies will be necessary to delineate the precise markers of neutrophil states and their associated plasticity in the human tumoral context.

Across multiple histologies, neutrophil gene signatures are associated with poor prognosis^42^, and the neutrophil-to-lymphocyte ratio (NLR) has been considered as a biomarker of poor survival outcomes in multiple therapies, including ICI ^43–45^. This is consistent with our data, wherein the neutrophil fraction (of CD45+ cells) is significantly higher in post-treatment non-responders when compared to post-treatment partial responders (Fig. 2f); this is further demonstrated via histology from a non-responding patient (MEL022) (Fig. 2g). However, calprotectin high, IL-8 high, and IFN-responsive neutrophils did not display a significantly larger representation in post-ICI non-responders compared to partial responders (Fig. S7). *CXCR2* and IL-8/*CXCL8* have both been linked elsewhere to ICI resistance as well as to the neutrophil N2 phenotype^46–48;^ however, as *CXCR2* (one of the two receptors for IL-8) was more highly expressed in calprotectin high neutrophils than in IL-8-high neutrophils (Fig. 2e), it may be that the protumor roles of neutrophils are not isolated to a single phenotype, but rather to a spectrum whose protumor effects are context-dependent.

Multiple mechanisms for the negative prognostic association of neutrophils have been proposed, including an association with tumor necrosis (itself linked to poor prognosis in multiple histologies)^49–59^, as well as the induction of angiogenesis, epithelial-mesenchymal transition (EMT), and neutrophil extracellular traps (NETosis)^60^.

Accordingly, we see a significant association (p=.0063) between samples’ neutrophil fraction (of CD45+ cells) and evidence of tumor necrosis on pathology reports (Fig 2h, Supplementary Table 1). Additionally, the presence of necrosis is particularly associated with the IL-8 high phenotype, although this association does not reach significance (Fig. 2h). Our data support the hypothesis that angiogenesis in the TME is promoted by IL-8 producing neutrophils, based on the significantly higher expression of genes in the hallmark angiogenesis gene module (see **Methods**) compared to the two other primary neutrophil phenotypes (Fig. 2i). We additionally see a significant correlation between patients’ IL-8 neutrophil fraction and the EMT module score of patients’ malignant cells (see **Methods**) (Fig. 2j). Lastly, genes associated with NETosis ^61^ were most highly expressed in calprotectin-high neutrophils but were also observed, albeit in decreasing amounts, amongst the IFN-responsive, and IL-8-high neutrophils, respectively (Fig. 2k). These collective findings suggest that neutrophils’ protumor function is multifaceted involving a spectrum of phenotypic states, and that a given neutrophil may play certain, though not all, protumor roles at a given time.

### Intracranial T cell phenotypic diversity consistent with that seen extracranially

We assessed the diversity of intracranial T cell phenotypes in the MBM-TME, and found that while our data recapitulated T cell phenotypic diversity observed extracranially, overall T cell infiltration did not reach statistical significance as a prognostic biomarker for ICI response in our cohort (Fig. 1e). In extracranial melanoma, both T cell infiltration and specific T cell phenotypes have been implicated as prognostic factors for response to ICI ^36, 62–65^. We therefore performed a focused analysis of only the flow cytometry- selected CD45+CD3+ cells within our scRNA-seq cohort (see **Methods**). Following a more stringent quality control screening via a cell complexity threshold of 2,000, a total of 2,974 cells were ultimately included for further analysis. Following unsupervised analysis, post-QC cells clustered into the following seven phenotypic populations, which we refer to as IFN-responsive, cycling, memory/naive, effector, exhausted, CD4/FOXP3, and NK/NKT cells (Fig. 3a,b). Cells within the “IFN-responsive” cluster were predominantly CD8+ T cells with upregulation of interferon response pathways (*IFIT1*, *IFIT2*, *IFIT3*, *ISG15*). The “cycling” population consisted of primarily CD8+ T cells with expression of canonical cycling genes (*MKI67*, *ZWINT*, *TOP2A*). Cells within the “memory” cluster included both CD8+ T and CD4+ T cells enriched for *IL7R*, and *CCR7*. “Effector” cells were marked by cytotoxic genes including *GNLY*, *GZMH*, *PRF1*, and *KLRG1,* and lower levels of exhaustion-associated genes than the “exhausted” cluster. “Exhausted” cells, meanwhile, expressed high levels *HAVCR2*, *PDCD1, CTLA4 and TIGIT*.

**Figure 3.**
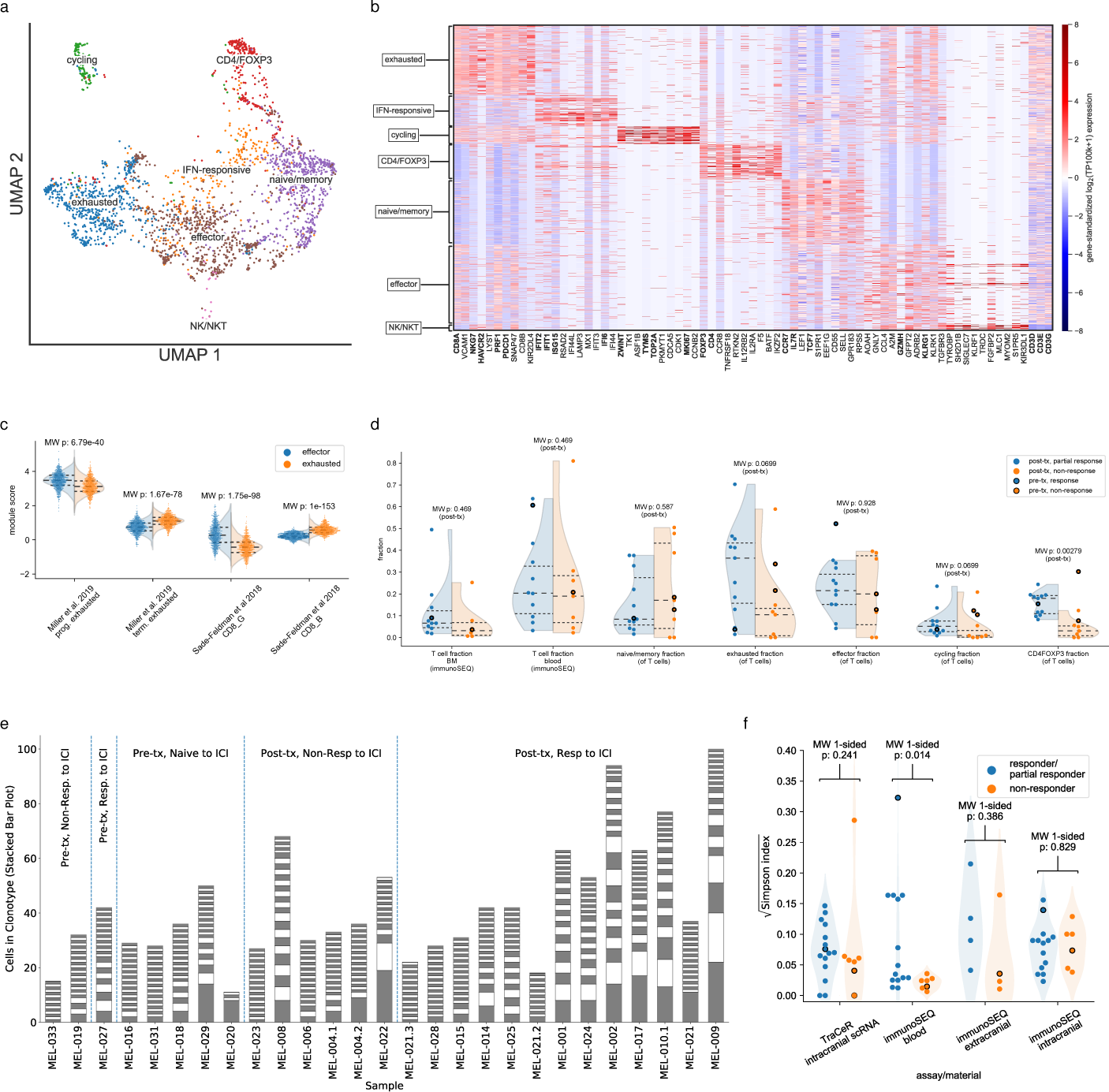
T cell phenotypic, clonotype heterogeneity, and corresponding association with response to immune checkpoint inhibition. a) UMAP of post-QC CD3+ (per flow cytometry) T cells. b) Heatmap standardized expression of top 10 marker genes for each cluster; key genes indicated in bold. c) Distribution (points indicating individual cells are overlaid on kernel density estimate of overall distribution) of module scores of previously discovered gene sets associated with progenitor vs terminal exhausted T cells (see **Methods**); Mann-Whitney U p-values for module scores between clusters indicated. d) Distribution (points indicating individual patients are overlaid on kernel density estimate of overall distribution of post-treatment patients) of T cell fraction (per the immunoSEQ assay) and phenotype fraction (of all post-QC T cells, per scRNA-seq data) across patients; pre-treatment patients noted in bold, Mann- Whitney U p-value comparing distribution of post-treatment partial vs non-responders indicated. e) Stacked bar plot for size of clonotypes identified via TraCeR, grouped by response to ICI, with size of bar indicating the number of cells in a clonotype (alternating colors used for visibility); samples with 10 or more successfully clonotyped cells included. f) Distribution (points indicating individual patients are overlaid on kernel density estimate of overall distribution of post-treatment patients) of Simpson indices according to TraCeR and immunoSEQ across patients; pre-treatment patients noted in bold, Mann-Whitney U p-value comparing distribution of post-treatment partial vs non- responders indicated. Samples are combined when multiple samples from a single patient were present.

The difference in phenotypes between the predominantly CD8+ “effector” and “exhausted” clusters are concordant with those observed in both murine models of melanoma ICI response ^65^, as well as those observed in clinical extracranial melanoma samples ^36^ (Fig. 3c), with the effector cluster corresponding to a progenitor exhausted phenotype/CD8_G, and the exhausted cluster corresponding to a terminally exhausted phenotype/CD8_B. However, while prior reports demonstrated that the ratio of “CD8_G” to “CD8_B” cells was associated with ICI response both before and following treatment ^36^, this association was not significant in our data; rather, the fractions of exhausted and cycling cells was associated (p=0.0699 for both) with partial vs non-response among the post-therapy samples (Fig. 3d). Among the post-therapy samples, only the CD4/FOXP3 fraction was significantly associated with partial over non-response (p=0.00279) (Fig. 3d). Neither intracranial nor blood T cell fraction (as measured by the immunoSEQ assay, see **Methods**) was associated with partial vs non-response post-therapy. We did, however, observe that the “effector” T cell fraction was highest in the pre-treatment responder sample in our cohort (MEL-027) (Fig. 3d), which is consistent with previous reports in the extracranial context ^36, 65^.

### Peripheral T cell clonal expansion associated with response to immune checkpoint blockade

Consistent with previous work^27, 66–70^ we observed that T cell clonal expansion was associated with response to ICI. From our cohort’s full-length transcript scRNA of freshly-resected metastases, we performed single cell T cell receptor (TCR) clonotyping via TraCeR^71^ (see **Methods**) to quantify T cell clonal expansion within the brain tumor microenvironment in addition to the genomic DNA based immunoSEQ assay (see **Methods**). Using TraCeR, we successfully clonotyped 2,110 pre-QC cells (with 1,371 clonotyped cells in the CD45+CD3+ post-QC group) from 31 unique samples (samples with greater than 10 clonotyped cells shown in Fig. 3e). We additionally assessed the presence of mucosal associated invariant T (MAIT) cells as well as invariant NKT (iNKT) cells (see **Methods**, Fig. S8), finding that putative MAIT cells are not significantly expanded relative to non-MAIT cells. We investigated the relationship between clonal expansion in both peripherally circulating and tumor infiltrating T cells using the Simpson index (a diversity-based measure of clonal expansion) across each compartment^72^, which was computed using both TraCeR- and immunoSEQ-derived TCR repertoires (see **Methods**). Using TraCeR results from within MBM, we found that patients who had demonstrated partial response to ICI with intracranial progression had a non-significant increase in their degree of clonal expansion compared to patients who were entirely non-responsive to ICI (p=0.241 via TraCeR, p=0.868 via immunoSEQ) (Fig. 3f). In contrast, sampling T cells from the blood demonstrated that clonal expansion tracked with partial response to ICI compared to the blood of patients who were non-responsive to ICI (P=0.014 via immunoSEQ) (Fig. 3f). These findings are consistent with previous reports suggesting that T cells from the peripheral circulation may be sampled as a biomarker of systemic response to ICI^73^.

### Intracranial clonally expanded T cells enriched for cells with exhausted phenotype

Given the association between clonal expansion and response to ICI, we assessed phenotypic correlates with clonal expansion. Of the 2,110 successfully clonotyped cells, 684 cells shared an α or β chain with one or more other T cells from the same patient, and were therefore regarded as being “detectably expanded” (448 of the 1,371 post-QC CD3+ T cells were detectably expanded). The remaining clonotyped cells we refer to as “non-detectably expanded” (Fig. 4a). Differential expression across samples of cells which were “detectably expanded” revealed enrichment of exhaustion- and effector-associated genes, including *HAVCR2*, *TIGIT, PRF1, NKG7*, and *GZMB*, in the detectably-expanded cells, whereas those cells which were clonotyped but not detectably expanded expressed higher levels of memory/naive-associated genes, including *CCR7* and *IL7R* (Fig. 4b). The majority of cells from two of the seven T cell clusters--”cycling” and “exhausted”--were detectably expanded (p=8.70e-7, 4.51e-59 respectively via Fisher’s exact test, see **Methods**) (Fig. 4c). By contrast, “memory” cells were most significantly enriched in the non-expanded set (p=7.83e-36 via Fisher’s exact test, φ coefficient=-0.3268, see **Methods**) (Fig. 4c). Thus, while clonal expansion is closely linked to response to ICI, we observed that clonally expanded cells at the site of the lesion were more likely to be exhausted.

**Figure 4.**
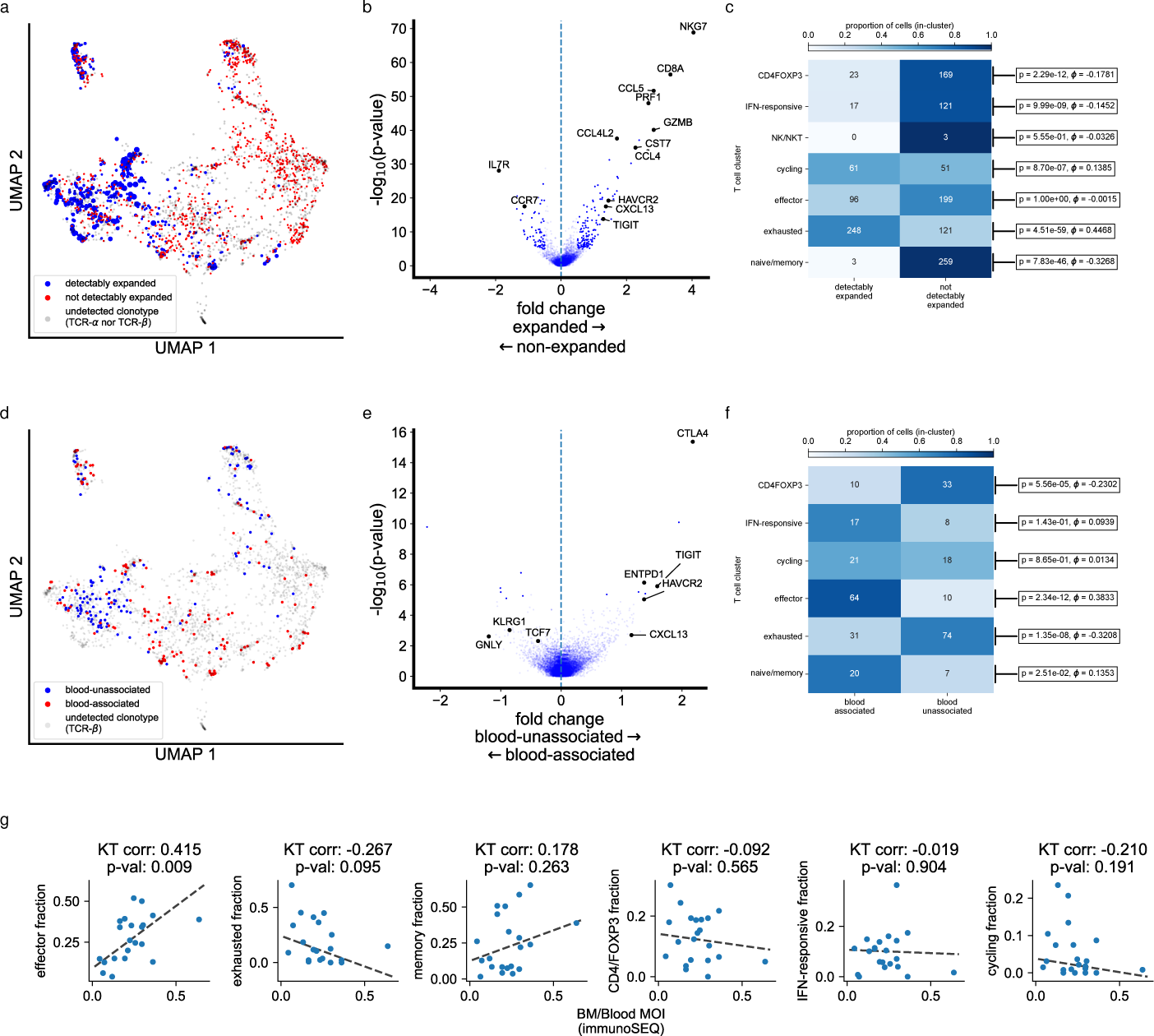
Association between T cell CDR3 and phenotype. a) UMAP (from Fig. 3a) indicating phenotypic distribution of detectably expanded T cells (blue), non-detectably expanded T cells (red), and non-clonotyped cells (gray); clone size of T cells belonging to detectably-expanded clone is proportional in size to marker area. b) Volcano plot of differentially expressed genes between detectably and non-detectably expanded cells (p-values calculated via Mann-Whitney U test, fold change described in **Methods**); key genes annotated. c) Heatmap of fraction of detectably and non-detectably expanded cells in each cluster. Colors indicate fraction in each cluster, on-block annotations indicate absolute number of cells in each stratum. P-values (Fisher’s exact test) and effect sizes (-coefficient) are annotated. d) UMAP (from Fig. 3a) indicating phenotypic φ distribution of blood-unassociated T cells (blue), blood-associated T cells (red), and non-clonotyped cells (gray). e) Volcano plot of differentially expressed genes between blood-unassociated and blood-associated T cells (p-values calculated via Mann- Whitney U test, fold change described in **Methods**); key genes annotated. f) Heatmap of fraction of blood-associated and blood-unassociated cells in each cluster. Colors indicate fraction in each cluster, on-block annotations indicate absolute number of cells in each stratum. P-values (Fisher’s exact test) and effect sizes (-coefficient) are φ annotated. g) Association between patient-averaged post-QC phenotype fraction and MOI (where patients contribute multiple samples, those samples are combined).Kendall-τ correlation and p-values (see **Methods**) are indicated. Theil-sen line of best fit (see **Methods**) indicated by dotted line.

### Intracranial T cells belonging to clonotypes detected in blood show reduced exhaustion

In order to investigate the relationship between clonally expanded T cells within the blood and tumor microenvironment, we performed TCR clonotyping of the TCR-β chain via the immunoSEQ assay of genomic DNA (gDNA) from peripheral blood (see **Methods**). T cell clones from the intracranial TME were subsequently matched (based on shared CDR3) against clones found in the blood. Intracranial T cell CDR3’s were therefore referred to as “blood-associated” T cell clones when they had a TraCeR- detected TCR-β CDR3 from the brain which were detected in the blood via the immunoSEQ assay. Those intracranial T cells with a TraCeR-detected TCR-β CDR3 which was not detected in the blood via immunoSEQ were denoted “blood- unassociated”. We observed a significant divergence in the distribution of “blood-associated” and “blood-unassociated T cells in our CD45+CD3+ UMAP (Fig. 4d). Differential expression of “blood-associated” and “blood-unassociated” T cells revealed upregulation of progenitor/effector-like genes (e.g. *TCF7, GNLY*, and *KLRG1*) in the former group, and exhaustion-related genes (*CTLA4*, *TIGIT, HAVCR2*) in the latter (Fig. 4e). Accordingly, we discovered a significant association between the “effector” population and blood-association (p=2.34e-12, Fisher’s exact test, φ oefficient=0.3833, see **Methods**) and a corresponding negative association with the “exhausted” population (p=1.35e-10, Fisher’s exact test, φ oefficient=-0.3208, see **Methods**) (Fig. 4f). Therefore, the Simpson index of the clonal repertoire measured in the periphery may be a more accurate measure of the reservoir of clonally expanding, non-terminally exhausted T cells at the time of sampling. These findings are consistent with findings in the context of extracranial melanoma^74, 75^. We validated this finding orthogonally on a patient-level basis by comparing phenotypic fraction with the Morisita Overlap Index (MOI, see **Methods**), a measure of TCR repertoire similarity between paired samples^76^. We observed that the fraction of CD45+CD3+ “effector” cells correlated significantly with the MOI between intracranial and blood-derived immunoSEQ TCR repertoires (p=.009, Kendall-tau correlation p-values, see **Methods**, Fig. 4g), suggesting that the MOI of blood and lesion-derived TCR repertoires can be used as an estimate of the effector-like T cell fraction within the lesion.

### T cell clones with private CDR3s are more likely to be exhausted and are associated with partial response to ICI

In order to explore the question of TCR tumor specificity in our cohort of intracranial melanoma metastases, we utilized a dataset of more than 500,000,000 clonotyped T cells across 1,486 samples from individuals with COVID-19 as a reference of presumably non-tumor specific clones (see **Methods**) and compared against our 2,110 clonotyped melanoma associated T cells^77^. A total of 68% of clonotyped post-QC CD45+CD3+ T cells from our melanoma brain metastasis cohort had CDR3s which were detected in the COVID-19 dataset suggesting that the majority of detectably expanded T cells within our cohort were not tumor specific. We subsequently categorized the melanoma T cell CDR3s as public, indicating that they were present in both our melanoma cohort and the COVID-19 data, or private, indicating that the T cell clones were solely identified in our cohort of MBM (Fig. 5a); as observed elsewhere^78–80^, these private CDR3s were significantly longer (two-sided Mann-Whitney U p-value=3.26e-11) than the public CDR3s (Fig. 5b). Upon further investigation of the phenotypic distribution amongst private and public clones, exhausted T cell clones were significantly associated with private CDR3 (p=0.00198) while an effector phenotype was primarily associated with public clones (p=0.0324) (Fig. 5c). These findings suggest that public T cell clones lack tumor specificity, and maintain an effector status without evidence of exhaustion. In contrast, private T cell clones appear tumor specific and may be driven to an exhausted phenotype by persistent antigenic stimulation.

**Figure 5.**
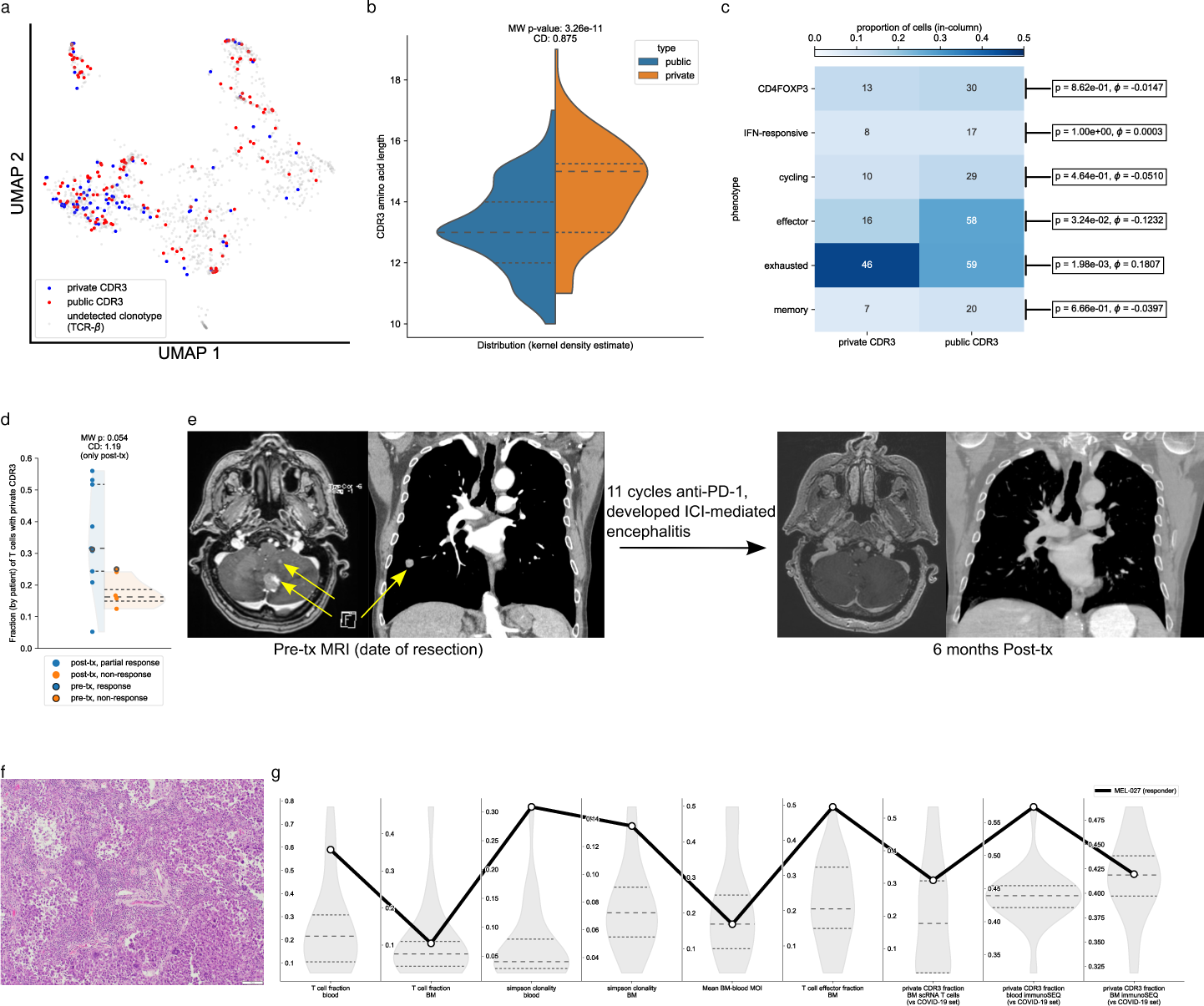
Association of clonotype privacy with phenotype and ICI response. a) UMAP (from Fig. 3a) indicating phenotypic distribution of T cells with private CDR3 (blue), public CDR3 (red), and non-clonotyped cells (gray). b) Distribution of CDR3 lengths in public and private clonotypes. Distribution is normalized kernel density estimate; quartiles indicated by dashed lines (median with longer dashed line). Mann- whitney p-value and Cohen’s D effect size annotated. c) Heatmap of fraction of cells with private and public CDR3 in each cluster. Colors indicate fraction in each cluster, on-block annotations indicate absolute number of cells in each stratum. P-values (Fisher’s exact test) and effect sizes (φ-coefficient) are annotated. d) Distribution (points φ indicating individual patients are overlaid on kernel density estimate of overall distribution of post-treatment patients) of fraction of T cells with private CDR3 across patients; pre-treatment patients noted in bold, Mann-Whitney U p-value comparing distribution of post-treatment partial vs non-responders indicated. e) T1 post-contrast MRI of the brain and lung from an ICI responsive individual (MEL027). Representative images of the brain and lung are included at the time of craniotomy for the large, symptomatic cerebellar metastasis (pre-ICI) and after six months of ICI administration (11 cycles of pembrolizumab) indicating resolution of intra- and extracranial disease. Yellow arrows mark regions of enhancement indicative of pre-treatment disease. f) Hematoxylin and eosin (H&E) staining of tumor section from MEL027, indicating large population of TILs. g) Multiple prognostic metrics across patients (see **Methods**); distribution for each metric, with upper, lower quartiles indicated by dotted lines, median by dashed line. Black line connects values for individual responding patient MEL027.

In order to examine the clinical relevance of this finding, we stratified patients by their clinical response to ICI and quantified the fraction of private T cell clones per patient. Within our cohort, we observed an association between a patient’s percentage of private CDR3 clones and overall response (p=0.054, two-sided Mann-Whitney U-test) (Fig. 5d).

Although all post-immunotherapy patients ultimately developed intracranial progression on immune checkpoint blockade which prompted surgical resection, MEL-027 was a treatment naive patient who went on to receive ICI and was the single patient within the cohort who was a true intracranial responder (Fig. 5e). With this unique clinical trajectory, we investigated the features of T cell infiltration and T cell clonal expansion within the blood and tumor microenvironment. The patient presented with multiple cerebellar metastases in addition to extracranial disease in the lung. Following resection of the dominant cerebellar lesion, pembrolizumab was initiated and resulted in both cranial and extracranial response (Fig. 5e). Histologic examination of the treatment naive cerebellar lesion demonstrated abundant tumor infiltrating lymphocytes (Fig. 5f).

This finding was corroborated with T cell fraction quantification from brain tumor and blood using the immunoSEQ platform (Fig. 5g, S9, see **Methods**). Across both sites, T cell fraction was above the median for the cohort (Fig. 5g, S9). Similarly, amongst our MBM cohort, MEL027 demonstrated the greatest degree of clonal expansion within the brain and blood while also harboring the greatest proportion of effector T cells within the brain (Fig. 5g, S9). Lastly, we explored the extent of T cell clonal overlap with clones from the COVID-19 dataset and observed that the fraction of private CDR3 clones for MEL027 in the blood was highest within the cohort and similarly elevated within the brain metastasis (Fig. 5g, S9). These findings are consistent with previously published reports that pre-treatment T cell infiltration is predictive of response to ICI while also integrating T cell phenotype, clonality, and tumor exclusivity.

## DISCUSSION

With the documented success of immune checkpoint blockade for extracranial disease and emerging data demonstrating intracranial response rates across multiple histologies, increasing attention has been placed on the tumor intrinsic and microenvironmental factors that portend a favorable response to ICI ^18^. Here, we utilize scRNA-seq from MBMs combined with TCR-seq from intracranial and extracranial samples to both characterize biomarkers of response to ICI, as well as to elucidate cross-compartmental correlates of malignant and immune phenotypes. We observe relationships between myeloid and malignant phenotypes, as well as relationships between T cells’ phenotype,TCR distribution, and diversity. These findings have implications for both the therapy and monitoring of intracranial disease.

Melanoma metastases have been significantly associated with increased levels of tumor infiltrating lymphocytes. Beyond T cell infiltration, however, increasing evidence suggests that the spatial distribution within the tumor, T cell phenotypic plasticity, and T cell clonality all impact anti-tumor immune mediated responses ^31, 69, 70, 81^. While single cell RNA sequencing has been used to characterize CD8+ T cell states in extracranial melanoma metastases, our cohort of MBM provides a unique opportunity to investigate the degree of phenotypic overlap within the brain metastatic TME. Prior studies identified naïve/memory, cytotoxic, and exhausted/dysfunctional T cells ^33, 34, 36^. We observe T cell transcriptional signatures that are concordant with those seen in extracranial melanoma^34–36^. While the fraction of effector-like T cells was elevated in the one pre-ICI responder in our cohort, neither this nor other CD8+ T cell phenotypes were significantly associated with partial vs non-response in the post-treatment samples.

T cell clonality is a proxy for antigen-driven T cell expansion and has been associated with clinical benefit across multiple tumor types and therapies^27, 67, 68^. Increasing efforts have attempted to understand the dynamics of T cell clonotype modulation, infiltration within the tumor microenvironment, and the relationship between T cell clones in the blood and tumor^69, 73, 82–84^. Within our cohort, clonal expansion in post-treatment blood, but not in intracranial lesions, was significantly associated with partial-response over non-response. This may be explained by the strong association between T cell clone size and exhaustion observed intracranially, wherein intracranial clonally expanded T cells have lost effector capacity due to persistent antigen stimulation, and therefore have reduced prognostic significance.

Our results provide critical context within the setting of CNS metastatic disease. Rather than representing a unique, immune isolated environment, our findings further support the concept that the CNS tumor microenvironment is immune-specialized rather than immune-privileged. In contrast, we observed unique phenotypic differences between detectably expanded and non-expanded T cell clones within the intracranial tumors.

While exhausted and cycling T cell clones were predominantly expanded, CD4/FOXP3 T cells, memory CD8 T cells, and IFN-responsive CD8 T cells were associated with non-clonally expanded T cells. These results are concordant with findings in both murine and clinical models of extracranial melanoma^74, 75^, where there is an association between a T cell clone’s phenotype and its shared presence in both the lesion and in the blood. This model of cross-talk between blood and extracranial tumors is similarly recapitulated within our cohort of melanoma brain metastases. With the finding of phenotypic divergence between blood-overlapping and tumor-exclusive T cell clones in MBM, this finding suggests a model where the CNS maintains features of the periphery with active cross-talk between the peripheral and intratumoral immune compartments with the blood acting as a reservoir of fresh, pre-antigen-stimulated T cells for MBMs, and that responding MBMs exist in an environment that is not fully immune-privileged^85^. Future investigation is needed to explore the process of intracranial T cell trafficking, tumor antigen exposure, and clonal replacement in brain metastases and cervical lymph nodes compared to extracranial sites of disease, the involvement of blood-brain-barrier disruption in this process, and how this process is further modulated by exposure to immune checkpoint blockade^21, 86–88^.

In addition to identifying individual T cell states, deciphering the nature of T cell tumor reactivity is an increasingly pressing challenge. The MBM-TME is populated both by T cell clones that are reactive to tumor antigens, as well as by bystander clones that are non-cancer-specific. Amongst our cohort of MBM, we observed a phenotypic difference between intracranial T cells with CDR3 regions that were patient-specific, and therefore assumed to be more likely specific to that patient’s tumor, in contrast to T cell clones with CDR3s found in public clonotype databases. Consistent with this finding, the greatest positive and negative associations with CDR3 privacy were observed in exhausted and effector cells respectively. The degree of CDR3 privacy was also associated with clinical benefit to ICI within the MBM context. Outside of MBM, increasing efforts have been made to understand features that identify tumor reactivity and delineate which phenotypic T cell states are most associated with response to ICI. Previously described features of tumor reactivity include a CD8+ T cell pre-dysfunctional state with expression of *CD39*, *CD103*, and *GZMK*; expression of immune checkpoints (*PDCD1*, *HAVCR2*, *LAG3*); oligoclonal TCR repertoire; and upregulation of immune activation markers (*TNFSF9* and *TNFRSF18*)^31, 89, 90^. Consistent with extracranial findings^31^, our data show that the majority of intracranial infiltrating T cells are not patient-specific. Collectively, these data suggest that T cell clonotype as well as phenotype should be jointly considered when using the T cell repertoire as a predictive biomarker in the context of MBMs.

In addition to exploring T cell features within the MBM-TME and the associated relationship with the periphery, our cohort provided a unique opportunity to explore the myeloid population in the MBM context. Dynamic features of myeloid phenotypic heterogeneity have been associated with both tumor-supportive and anti-tumor properties ^91^, however, their role in the CNS is not fully understood. Recent studies have utilized single-cell genomics of the monocyte population within the context of normal CNS development, CNS insult, and malignancy^37, 92–94^. Consistent with work from Klemm et al.^94^ and Friebel et al.^93^, we observed multiple monocyte phenotypes within MBM, including *TREM2/APOE* expressing reactive microglia^37^. *TREM2* expression has been inversely correlated with overall survival across multiple histologies^95, 96^. Utilizing melanoma scRNA-seq datasets, Xiong et al.^97^ has linked *TREM2*-expressing macrophages with complement activation, tumor-associated macrophage polarization, and ICI resistance^97^--a finding similarly observed by Molgora et al. within the context of *in vivo* sarcoma models^95^. While we observed *TREM2*+ reactive microglia in too few samples to evaluate whether they play a significant role in the context of MBM response to ICI (Fig. S4a), further work is warranted to explore (1) the role of these cells in modulating the immune microenvironment within the context of melanoma brain metastases, (2) the relative contribution of *TREM2*-expressing monocyte in the intracranial and extracranial compartment, and (3) the potential of targeting this population to enhance immune checkpoint blockade for CNS metastatic disease.

Our analysis of the myeloid compartment provided unique insights into neutrophil heterogeneity within the MBM-TME, which recommend further studies with larger patient cohorts and myeloid cell numbers. We identified a significant association between an IL-8 expressing neutrophil subset and EMT transition in neutrophils. While this neutrophil subset has been previously identified, and may correspond to the N2 subset described in other contexts, they have not, to our knowledge, been previously observed in MBMs. Serum IL-8 levels have previously been reported to be associated with worse prognosis and reduced clinical benefit of ICIs^48, 98^. Due to the significantly higher infiltration of neutrophils in intracranial relative to extracranial tumors^99^, it is possible that these IL-8 expressing neutrophils play an even greater role in determining intracranial prognosis and ICI responsiveness than they do extracranially. We do note certain discrepancies between previously described N1 and N2 markers and markers observed in our scRNA-seq data, suggesting that the precise markers of different neutrophil phenotypes--and their association with neutrophil-mediated phenomena including angiogenesis and NETosis--are context-dependent. Further work is needed to dissect this cell population’s phenotypic plasticity, link cell state with tumor supportive or suppressive roles, investigate their relative distribution across histologies, and study their differential role within intra- and extracranial metastases.

The field of immune checkpoint blockade has revolutionized the treatment of metastatic melanoma and is increasingly being applied to an array of other histologies with encouraging results. Unfortunately, the CNS is frequently a site of disease progression and ultimate patient mortality. While recent clinical trials exploring ICI for melanoma brain metastases have demonstrated encouraging results, a significant number of patients continue to experience CNS progression. The results of our study provide unique insights into the relationship between features of the tumor microenvironment of the brain and previous findings within the extracranial compartment. Moreover, as many myeloid phenotypes are specific to the CNS or elevated intracranially^99^, they may represent particularly attractive targets for increasing intracranial ICI efficacy. For example, the role of unique myeloid populations including *TREM2*/*APOE* expressing monocytes and *CXCL8*/*VEGFA* expressing neutrophils appear to be linked to tumoral plasticity and shifts in ICI responsiveness. Further investigation will be needed to explore the presence and longitudinal dynamics of these populations in the cranial and extracranial compartments for metastatic melanoma.

Within the T cell compartment, single cell immune profiling with associated TCR clonotyping provided clues regarding responsiveness in the brain metastases setting. We find that multiple factors--T cell fraction, phenotype, and clonotype--play roles in determining intracranial MBM response to ICI, and recommend that future studies jointly consider these factors whenever possible. Our cohort also provided the unique opportunity to explore the TME of a treatment-naive individual (MEL-027) who went on to respond to ICI. From a histologic perspective, the tumor demonstrated robust lymphocyte infiltration throughout the tumor suggestive of an immunologically “hot” tumor. Additionally, features of the blood compartment were reflective of a baseline pro-inflammatory state with elevated T cell fraction, simpson index, and fraction of private T cell clonotypes. These features of the periphery were recapitulated within the brain metastasis with an elevated T cell fraction, abundance of effector T cell clones, robust clonal expansion, and concomitant elevation of private T cell clones within the tumor.

While our study provides insights into features defining the MBM-TME and potential factors that reflect response to ICI, our results have several limitations. Our cohort reflects a relatively small population of patients with both diverse treatment courses and varied responses to therapy across multiple sites of disease. Additionally, we were limited by a lack of matched pre- and post-ICI brain metastasis samples--a challenge inherent with this patient population. While we were able to analyze a single treatment naive patient who responded to ICI, a larger cohort of treatment naive individuals who subsequently are treated with ICI is needed to extend the applicability of those findings. We were similarly limited by the ability to sample patient matched cranial and extracranial tumors in order to precisely decipher at a single cell resolution the intratumoral features that are unique to the MBM-TME.

Nevertheless, the implications of this unique cohort of patients provide intracranial context within the broader context of immunotherapy for metastatic melanoma. Our collective results emphasize (1) the critical role of T cell mediated response in the setting of ICI for MBM, (2) elucidate the relationship between T cells within the blood and intratumoral compartment, and (3) demonstrate that blood provides insights into ICI response not only for extracranial disease, but also disease within the brain. Moreover, the relationship between T cell clonal expansion and phenotypic states in the blood and brain has the potential of providing critical insight into the intracranial TME, which may be clinically advantageous when acquisition of brain tumor tissue is limited. Lastly, future work using larger cohorts of human specimens and murine models will be needed to fully understand the intracranial features of the tumor microenvironment that attenuate these initially robust intracranial responses.

## METHODS

### Collection of fresh tissue for scRNA-seq

This study was conducted in accordance with recognized ethical guidelines and patient samples were collected under the Dana-Farber Cancer Institute’s (DFCI) Institutional Review Board (IRB) approved protocol 10-417, Tissue Bank for Neurological Disorders. All participants signed informed consent prior to collection of specimens. Thirty-two sequentially collected MBM from 27 patients were obtained immediately following craniotomy. Tumors were classified according to prior exposure and therapeutic response to ICI. Patient intracranial and extracranial clinical responses were categorized as Responder, Partial-Responder, and Non-Responder after review of clinical history and imaging with board certified medical oncologist, neuro-oncologist, and radiologist. An overview of the clinical features of the cohort are shown in Supplementary Table 1. Collection of solid tumor tissue and blood samples was performed to reduce the time between collection and processing. Approximately 20mL of blood is collected at any time point during the patient’s surgery with coordination and permission of the anesthesiology team. Blood is collected in two 10mL EDTA tubes that we provide. Immediately after collection, a portion of the blood is used to separate plasma, and another portion of whole blood is saved for eventual gDNA extraction. Solid tumor tissue is collected via the coordination of lab technicians with the participant’s surgical team. Lab technicians collect tissue directly from the operating room and bring it to the pathology team to approve a portion of the tissue to be used for research. The tissue is then immediately brought back to the lab to begin the dissociation workflow (described below).

### Tumor dissociation workflow

After collection, the specimen was transferred to a sterile petri dish and mechanically dissociated with a scalpel. The dissociation mixture used 4mL of preheated Buffer X, 40uL of Buffer Y, 50uL buffer N and 20uL of enzyme A from the Miltenyi Biotec Brain Tumor dissociation kit, a papain based dissociation kit (Miltenyi Biotec, cat. no. 130-095-942). The tumor was combined with the digestion buffer and incubated on a rotator at 37°C for 30 minutes. The tissue was then resuspended thoroughly via pipette, passed through a 100 micron cell strainer and spun down at 188g for 3 minutes. The pellet was washed with 5mL of HBSS with calcium and magnesium and ultimately resuspended in 90uL of PBS + 1% BSA.

### Flow cytometry analysis

The cell mixture was stained with 10uL of CD45-Vio-blue (Miltenyi, cat. no. 130-113-122) and 10 uL of CD3-PE (Miltenyi, cat. no. 130-113-139), then incubated on ice for 20 minutes.

Following this incubation,1.5uL Calcein-AM live cell stain (Thermo Fisher, cat. no. C1430) and 0.5 uL To-Pro-3 Iodide dead cell stain (Thermo Fisher, cat. no. T3605) were added to the cell suspension. The same cells were used for unstained controls to adjust gating. Following dissociation and staining, single cells were purified by fluorescence-activated cell sorting (FACS) with the BDFACSAria Fusion instrument, which employs five lasers (405 nm, 488 nm, 640 nm, 355 nm and 561 nm). Cells were sorted into fully skirted 96-well plates with 10ul of buffer TCL+1% beta-mercaptoethanol. Set up for each experiment used a 100-um nozzle at 20 psi and 31 kHz. Prior to fluorescence gating, forward scatter (FSC-A) and side scatter (SSC-A) were used to identify granularity and ensure to sort only singlets. We then used fluorescence staining to gate conservatively and specifically for live cell populations within the CD45+, CD45-, and/or CD45+ CD3+ cells clusters. The sorted plates were then spun down at 188g for 1 minute, flash frozen at -80°C and stored for future scRNA sequencing. Flow sorting analysis was completed with the FACSDiva (v. 8.0.1) and FlowJo (v. 10).

### scRNA-seq

Single cell RNA sequencing was performed using the Smart-Seq 2 protocol to create an 8 plate (4 CD45-, 2 CD45+ and 2 CD3+) cDNA library for each patient tumor sequenced. RNA isolation is first completed by resuspending each well with 22 uL of RNAclean XP beads (Beckman, cat. no. A63987) and transferring the mixture to a demi-skirted 96 well PCR plate. The cells were incubated at room temperature in the bead suspension, then transferred to a magnetic plate and washed twice with 80% ethanol. Reverse transcription was performed using Maxima H minus reverse transcriptase (Thermo Fisher, cat. no. EP0753) and 10 uM of TSO oligonucleotides (Exiqon), and incubated at 42C for 90 minutes then 10 cycles of (50°C for 2 min, 42°C for 2min) then heat inactivation at 70°C for 15 min. Full length cDNA amplification with Hi-Fi Hotstart Readymix (Roche, cat. no. 07959079001) was completed with a 98°C incubation then 21 cycles of (98°C for 15 sec, 67°C for 20 sec, 72°C for 6 min) and a final extension at 72°C for 5 min.

Amplified cDNA was purified with AmpureXP beads (Beckman, cat. no. A63881) and cleaned with two 80% ethanol washes on the plate magnet. An Agilent BioAnalyzer high sensitivity DNA chip (Agilent Technologies, cat. no. 5067-4626) was used to ensure proper distribution and fragment length of the cDNA library. cDNA library concentration was measured using the Qubit dsDNA HS assay (Thermo Fisher, cat. no. Q32854) according to the manufacturer’s protocol and values were read on a microplate reader. Using the Qubit assay concentration values, all library wells were diluted to 2 ng with water in order to proceed to library preparation. The Nextera XT Library Prep Kit (Illumina, cat. No. FC-131-1096) protocol was used for fragmentation and unique barcoding of the cDNA libraries. For each plate, 1.5 uL per well was pooled and purified with the Ampure XP beads (0.9X the amount of sample) and two 80% ethanol washes. The cDNA library pools were run on BioAnalyzer high sensitivity DNA chip to calculate base pair size, followed by a Qubit assay on the Qubit Fluorometer to determine cDNA concentration; ultimately, these values were used to dilute the samples to 2 nm with water. 5 uL of eight pools were multiplexed together for sequencing. Samples were sequenced using a NextSeq 500/550 instrument (Illumina) with the Nextseq 500 High Output v2.5 75 cycle kit (Illumina, cat. no. 20024906).

### gDNA extraction from blood

500uL of whole blood was used to extract gDNA using the Qiagen blood and tissue kit (Qiagen, cat. no. 69506). Blood was processed according to the manufacturer’s guidelines no later than two weeks post collection, while being stored at 4°C, and after extraction gDNA was stored at - 20°C until sequencing.

### gDNA extractions from fresh frozen tissue (FF)

Between 25-35mg of fresh frozen tissue, stored at -80°C, was first mechanically dissociated using an RNAse free homogenizer until the tissue was completely dissociated into solution. gDNA was then extracted using the Qiagen AllPrep DNA/RNAmiRNA Universal kit (Qiagen, cat. no. 80224) according to the manufacturer’s guidelines. Samples were stored at -20°C until sequencing.

### gDNA extractions of tissue from formalin-fixed paraffin-embedded (FFPE) slides

One slide from each case was first stained with hematoxylin and eosin then evaluated by a collaborating pathologist to determine the location of tumor tissue on the paraffin slide. Slides were then scraped to strategically collect the tumor tissue into 1.5 mL eppendorf tubes containing deparaffinization solution. gDNA was then extracted using the Qiagen QIAmp DNA FFPE Tissue kit (Qiagen, cat. no. 56404) according to the manufacturer’s guidelines. Samples were stored at -20°C until sequencing.

### Quantification of DNA

The Pico green assay (invitrogen, cat. no. P11496) was used according to the manufacturer’s protocol to quantify the concentration of DNA extracted from FFPE, fresh frozen tissue, and blood.

### Whole-exome Sequencing

Whole-exome sequencing was conducted on extracted DNA on an Illumina HiSeq platform at the Broad Institute. Library prep and exome enrichment was performed using either the Illumina Content Exome (ICE) or TWIST Somatic exome v6 platforms. Standard Coverage ICE exomes were 80% of targets at 20X, Deep Coverage ICE exomes were 85% of targets at 50X, and TWIST Somatic v6 exomes were 85% of targets at 100X. Sequencing for ICE was then performed using 76 bp runs with an 8 base index sequencing read on Illumina HiSeq RTA v1.18.64 or later. Sequencing for TWIST used 151 bp runs on Illumina’s NovaSeq S4.

### DNA-based TCR-β sequencing

The Adaptive Biotechnologies immunoSEQ human T-cell receptor beta (hsTCRB) v4b kit was used to identify and quantify the frequency of specific T-cell clones in extracted DNA. DNA that was extracted from FF, FFPE-preserved tissue, and blood samples, as explained earlier, was quantified with the PICO assay to use for TCR sequencing. DNA from FF samples were diluted to 83 ng/ul and run in duplicates, gDNA from blood samples were diluted to 44 ng/ul and also run in duplicates, while all extracted DNA from FFPE samples was divided and run in quadruplicates. A negative control is also included and run in parallel with the samples. After the two PCR amplification steps, given in the manufacturer’s protocol, the multiplexed sample was run on an agilent BioAnalyzer high sensitivity DNA chip (Agilent Technologies, cat. no. 5067- 4626) and a KAPA Library qPCR quantification kit (Roche, cat. no. 07960140001). Sequencing was performed on an Illumina NextSeq 500/550 instrument (Illumina) using the Nextseq 500 High Output v2.5 150 cycle kit (Illumina, cat. no. 20024907). DNA-based TCR- β rearrangement repertoires, as well as sample T cell fractions, were then obtained via the immunoSEQ Analyzer pipeline (Adaptive Biotechnologies).

### Reconstruction of T cell receptor (TCR) from scRNA-seq data

TCR α and β chains were reconstructed using TraCeR (https://github.com/Teichlab/tracer) .

### Analysis of TCR data

The Simpson index was computed without replacement for TraCeR samples according to the formula

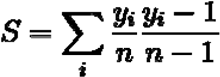

The Simpson index was computed with replacement for immunoSEQ samples according to the

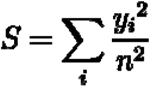

formula where is the total number cells in a sample, is the number of cells in clonotype .

The Morisita-Horn Overlap Index between samples and is computed according to the formula

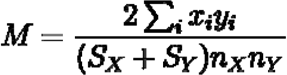

For Simpson indices, for samples and respectively, computed without replacement;, are total cell counts for samples, respectively, with, representing the number of cells in clonotype

### Alignment & Pre-processing of scRNA-Seq data

Illumina sequencing outputs were demultiplexed via the bcl2fastq2 program, v2.20.0.422. Generated fastq files were aligned and corrected for PCR bias using the RSEM program^100^ using version 8 of the smartseq2 workflow on Terra (app.terra.bio) provided by Cumulus^101^. Transcripts were aligned using Bowtie 2^102^ to human genome GRCh38, with gene annotation generated using human Ensembl 93 GTF.

### Unsupervised transcriptomic analysis

All analyses were done using the panopticon package^103^. Resultant count matrices were combined and normalized via panopticon.preprocessing.generate_count_matrices. Principal components analysis was performed via the command panopticon.analysis.generate_incremental_pca. UMAP embedding was performed with the command panopticon.analysis.generate_embedding with default parameters. Clustering was performed via the command panopticon.analysis.generate_clustering, which performs multiple rounds of agglomerative clustering with a correlation metric, with the number of clusters selected via a silhouette score. Feature selection (first 10 principal components) is recomputed at each subsequent round, wherein clustering within previously identified clusters is performed. Source code for the panopticon package can be found at https://github.com/scyrusm/panopticon, with additional documentation at https://panopticon-single-cell.readthedocs.io/en/latest/.

### Cell quality control

Cells were initially filtered based on a minimum unique gene count of 1000. Clustering was then performed as described above. These clusters were then manually reviewed to assess whether they showed signs of contamination, or of being doublets. Results of this manual review, and justification for the inclusion or removal of cells, are given in Supplementary Table 2.

### Detecting malignant cells using inferred copy number variations

Inferred copy number profiles were performed within cells from a single patient taken all FACS gating categories (CD45-, CD45+, CD3+) according to the procedure in Tirosh et al.^34^. These copy number profiles were than projected onto their first principal component (within a group of cells from a single patient). The quantile of cells’ loading onto this component was denoted the “malignancy score,” with the loading sign-adjusted such that the CD45- cells (per FACS) had a greater such mean score than the grouped CD45+/CD3+ cells. This score was used as a factor when considering cell quality control above (see Supplementary Table 2). Code for this procedure is implemented in panopticon.analysis.generate_malignancy_score.

### Differential expression and gene expression plots

DIfferential expression between sets was computed with the Mann-Whitney U test, as implemented in panopticon.analysis.cluster_differential_expression, between groups using log2(TP100k+1) gene expression values. Dot plots (Fig 2e,k) were computed via the function panopticon.visualization.plot_dotmap.

### Gene expression Signatures

Module scores were as originally used in Tirosh et al.^34^, implemented in the panopticon package as panopticon.analysis.generate_masked_module_score, over the set of cells as described in the text. The following MSigDB v7.4 signatures^104, 105^:

- Epithelial mesenchymal transition score (EMT): https://www.gsea-msigdb.org/gsea/msigdb/cards/HALLMARK_EPITHELIAL_MESENCHYMAL_TRANSITI ON.html
- Interferon gamma response: https://www.gsea-msigdb.org/gsea/msigdb/cards/HALLMARK_INTERFERON_GAMMA_RESPONSE.html Eosinophil markers (NAKAJIMA_EOSINOPHIL): https://www.gsea-msigdb.org/gsea/msigdb/cards/NAKAJIMA_EOSINOPHIL.html
- Primary (azurophilic) granule genes (GO_AZUROPHIL_GRANULE): https://www.gsea-msigdb.org/gsea/msigdb/cards/GOCC_AZUROPHIL_GRANULE.html
- Secondary (specific) granule genes (GO_SPECIFIC_GRANULE): https://www.gsea-msigdb.org/gsea/msigdb/cards/GOCC_SPECIFIC_GRANULE.html Tertiary granule genes (GO_TERTIARY_GRANULE): https://www.gsea-msigdb.org/gsea/msigdb/cards/GOCC_TERTIARY_GRANULE.html
- Angiogenesis: https://www.gsea-msigdb.org/gsea/msigdb/cards/HALLMARK_ANGIOGENESIS.html
- We additionally used signatures from Sade-Feldman et al. 2018 (CD8_G, CD8_B)^36^, Miller et al. 2019 (progenitor exhausted, terminally exhausted)^65^, and Li et al. 2019 (cell stress signatures)^33^. These signatures are given in Supplementary Table 3.

### Statistical analyses

All calculations were performed using python v3.7.4. The following functions and associated p-values were used from the scipy package, v1.5.4: Mann-Whitney U (scipy.stats.mannwhitneyu), Kendall-τ (scipy.stats.kendalltau), Fisher’s exact test (scipy.stats.fisher_exact), Theil-Sen slopes (scipy.stats.theilslopes). All p-value tests were two-sided unless otherwise noted. The panopticon package^103^ v0.1.1 was used throughout; Cohen’s d and phi-coefficient effect sizes were computed via the panopticon.utilities.cohends and panopticon.utilities.phi_coefficient functions.

Fold changes in Fig. 4b,e were computed as the difference in the means of log2(TP100k+1) expressions between two groups. Fisher’s exact tests for each row in Figs. 4c,f were computed according to the following contingency table

|  |  |
|---|---|
| a | b |
| c | d |

where a, b represent the number of cells of the phenotype in question (CD4/FOXP3, IFN-responsive, NK/NKT, cycling, effector, exhausted, naive/memory) which are detectably expanded/not-expanded, respectively (for Fig 4c) or blood-associated/non-associated, respectively (for Fig 4f). Table elements c, d represent the sum of cells belonging to all other phenotypes which are detectably expanded/not-expanded, respectively (for Fig 4c) or blood-associated/non-associated, respectively (for Fig 4f). The Fisher’s exact test p-value is

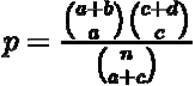

computed in the usual way, namely where is the binomial coefficient. The same contingency table is used to compute the phi coefficient via the usual formula:

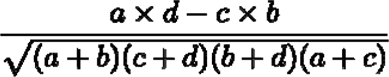

Throughout, kernel density estimate plots were computed via the seaborn.violinplot package using seaborn v0.11.0, with the argument “cut=0, inner=’quartile’” all other parameters default.

### Analysis of putative iNKT, MAIT cells

Putative iNKT cells, MAIT cells were classified according to known TCR-α V/J allele combinations, or TCR-β V alleles that have been associated with these cells, according to Mori et al. (see Table 3)^106^.

### Assessing CDR3 public/private status

A large cohort of TCR-β repertoires was obtained from the immuneCODE dataset (https://clients.adaptivebiotech.com/pub/covid-2020)77. TCR-β CDR3s detected via TraCeR were classified as being private or public based on whether they were detected at any level in the immuneCODE dataset.

## DATA AVAILABILITY

The genes-by-cells matrix and associated metadata for the current study, including all UMAPs for plots in this manuscript, are available via the Broad single-cell portal: https://singlecell.broadinstitute.org/single_cell/study/SCP1493/microenvironmental-correlates-of-immune-checkpoint-inhibitor-response-in-human-melanoma-brain-metastases-revealed-by-t-cell-receptor-and-single-cell-rna-sequencing. Raw data, including fastq files from both scRNA-seq and whole exome sequencing, are available on dbGAP (accession pending). TCR-seq data generated through the immunoSEQ assay is available through the immuneACCESS portal (link pending).

## CODE AVAILABILITY

Panopticon v0.1.1 has been made publicly available (https://github.com/scyrusm/panopticon/). Notebooks used for figure creation available upon reasonable request.

## ACKNOWLEDGEMENTS

Funding for this research was provided by Merck Sharp & Dohme Corp., a subsidiary of Merck & Co., Inc., Kenilworth, NJ, USA; Damon Runyon Cancer Research Foundation; Melanoma Research Alliance; Breast Cancer Research Foundation; and 1R01CA244975-01. PKB and SLC were supported by the NIH (5R01CA227156-02, 1R21CA220253-0A01 and 1R01CA244975-01). PKB was also supported by Susan G. Komen. SLC was also supported by the Wong Family Award, the DF/HCC Lung Cancer Program 25Developmental Research Project Award in Lung Cancer Research, and by Dana-Farber Institutional Research Support. CAB was supported by an NIH/NINDS R25, the American Brain Tumor Association, and AANS/CNS Joint Section on Tumors. BI was supported by NIH R37CA258829 and R21CA263381, and the Burroughs Wellcome Fund Career Award for Medical Scientists. BCM is supported by the National Cancer Institute of the National Institutes of Health under 1K08CA248960 and the Wong Family Award. MAD is supported by the Dr. Miriam and Sheldon G. Adelson Medical Research Foundation, the AIM at Melanoma Foundation, the NIH/NCI (1 P50 CA221703-02 and 1U54CA224070-03), the American Cancer Society and the Melanoma Research Alliance, Cancer Fighters of Houston, the Anne and John Mendelsohn Chair for Cancer Research, and philanthropic contributions to the Melanoma Moon Shots Program of MD Anderson.

## AUTHOR CONTRIBUTIONS

CAB, SLC, RJS and PKB designed the study. CAB, JHS, AEK, MRS, MdS, AD, JLM, NN, JML, AK, MW, BS, and CMG collected clinical patient data. DTF, GMB, MML, MF, BVN, WTC, BSC, and DPC assisted with tissue acquisition. CAB, JHS, MRS, AEK, JLM, NN, MW, MML, MF, NW, EEG, RJS, and PKB reviewed clinical, radiographic, and pathology results. CAB, NN, JHS, MdS, MS, AD, AWN, AJ, BMK, and JLM collected and processed samples. CAB and JHS performed single-cell sequencing experiments. MLS, BI, MD, CH, KP, BM, OS, AS, and MS provided experimental and analytical support. ML and SCM analyzed scRNA-Seq data under the guidance of SLC and PKB. CAB, SCM, SLC, and PKB wrote the manuscript with input from all authors.

## SUPPLEMENTARY FIGURES

**Supplementary Figure 1.**
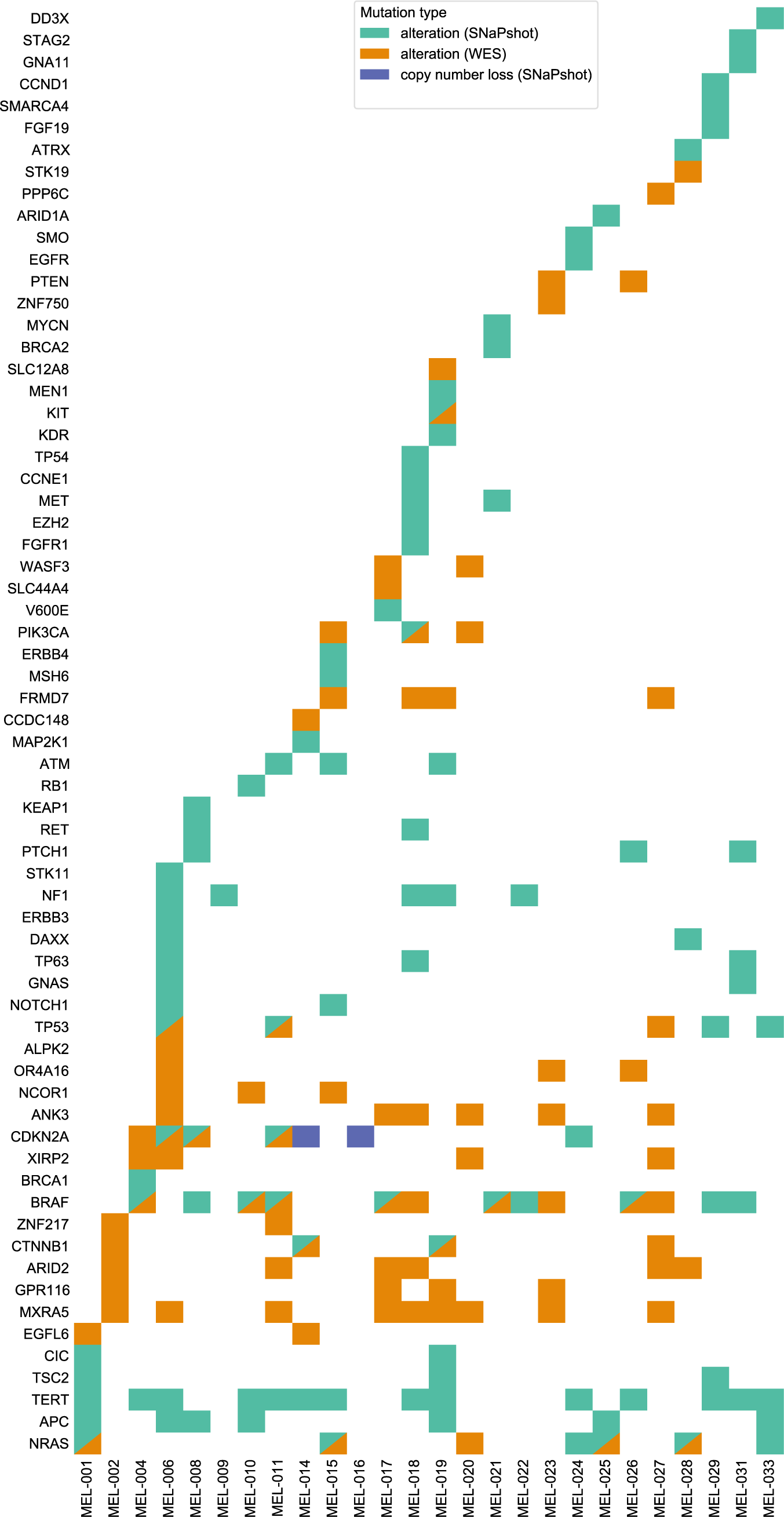
Comut plot of melanoma driver alterations, per WES, SNaPshot. Presence of alterations, detected through SNaPshot or whole exome sequencing (WES). Type of alteration, where relevant, indicated in legend.

**Supplementary Figure 2.**
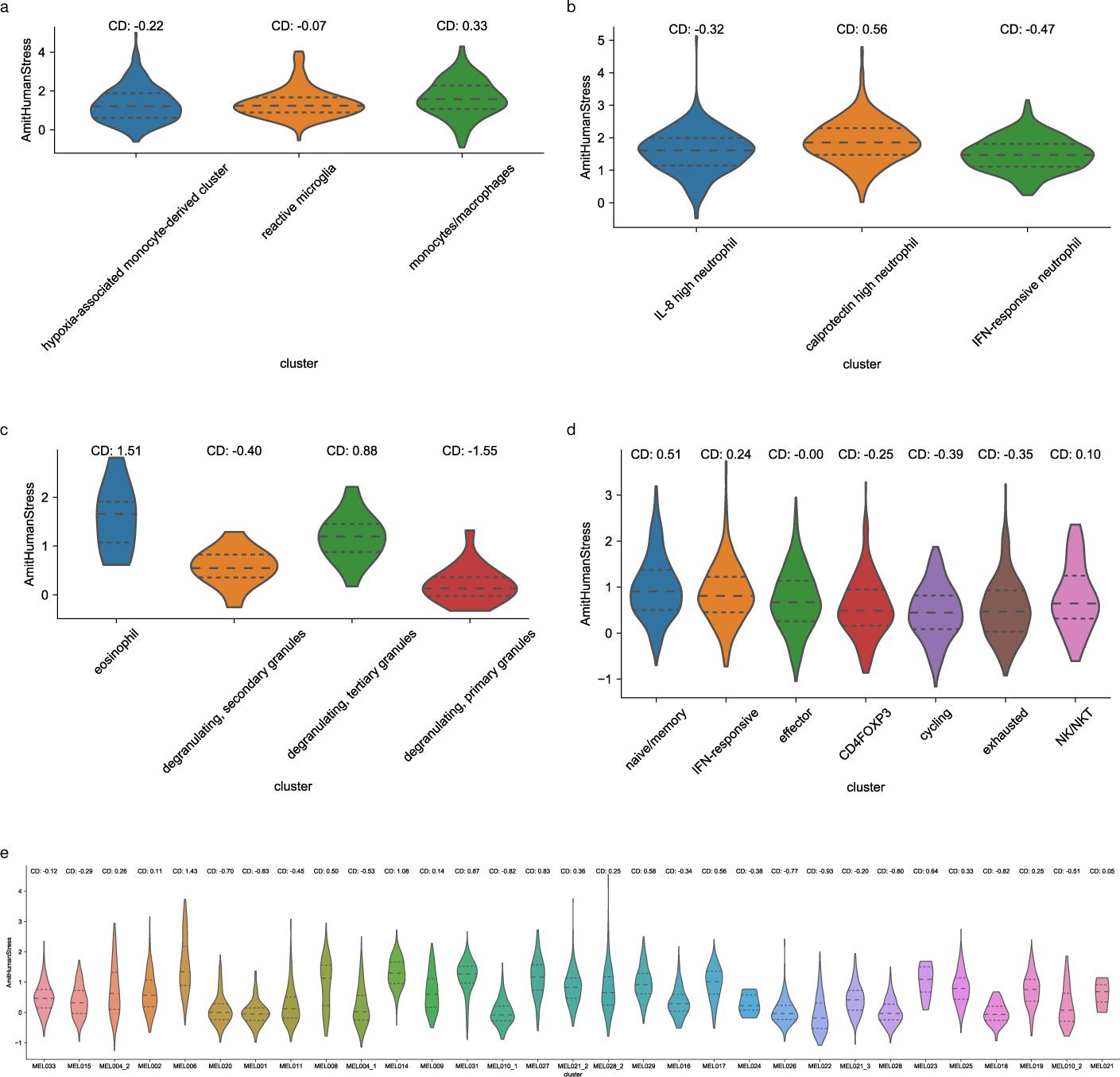
Cell stress signatures across cell types. Violin plots with dashed quartile lines indicating module scores of cell stress signature (see **Methods**) for a) monocyte-derived/microglial cells, b) non-degranulating neutrophils, c) degranulating neutrophils, eosinophils, d) post-QC T cells, and d) malignant cells, grouped by sample.

**Supplementary Figure 3.**
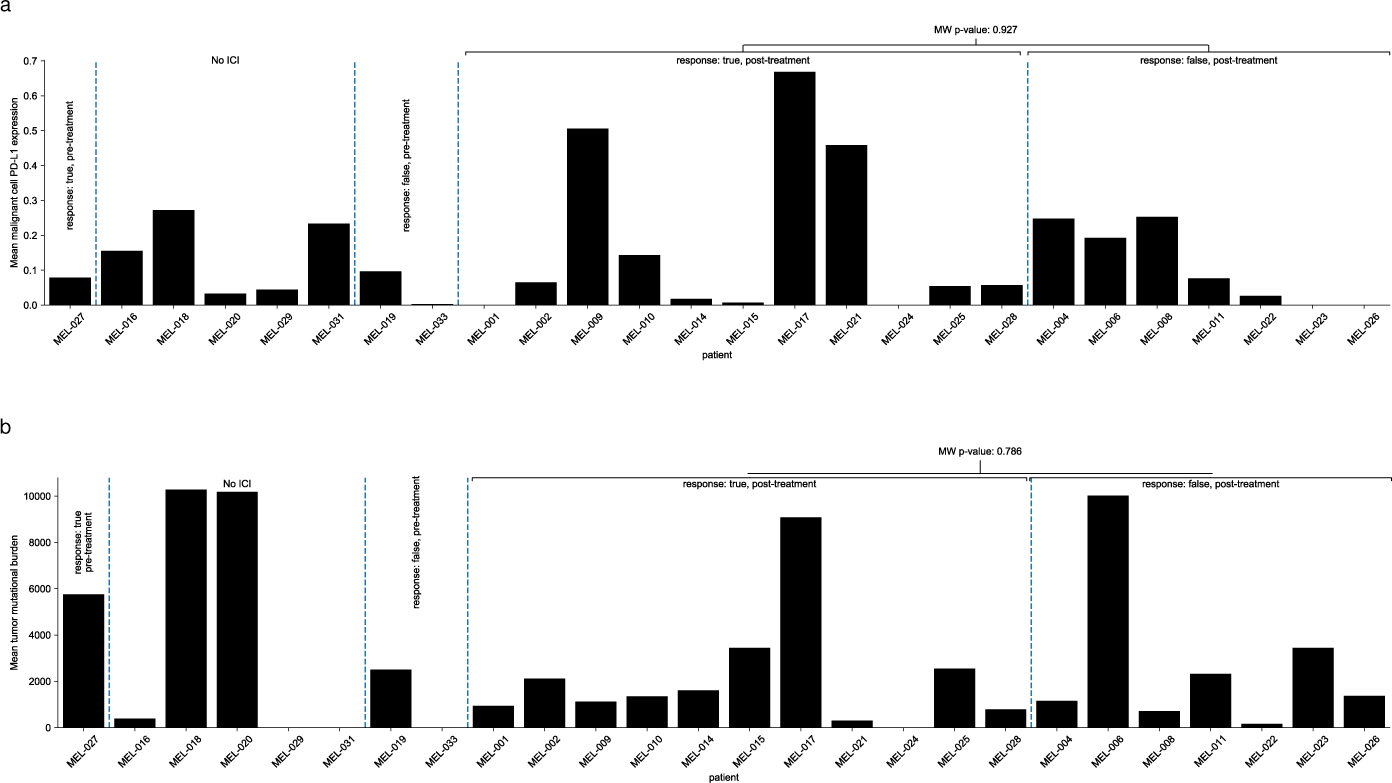
Malignant cell PD-L1, TMB levels across patients. a) Mean PD-L1 expression among malignant cells across patients (where multiple samples from a single patient exist, expression levels are averaged). Mann-whitney p-value for PD-L1 levels between post-treatment partial-responding and non-responding patients indicated. b) Tumor mutational burden (TMB) levels across samples. Mann-whitney p-value for TMB levels between post-treatment partial-responding and non-responding patients indicated.

**Supplementary Figure 4.**
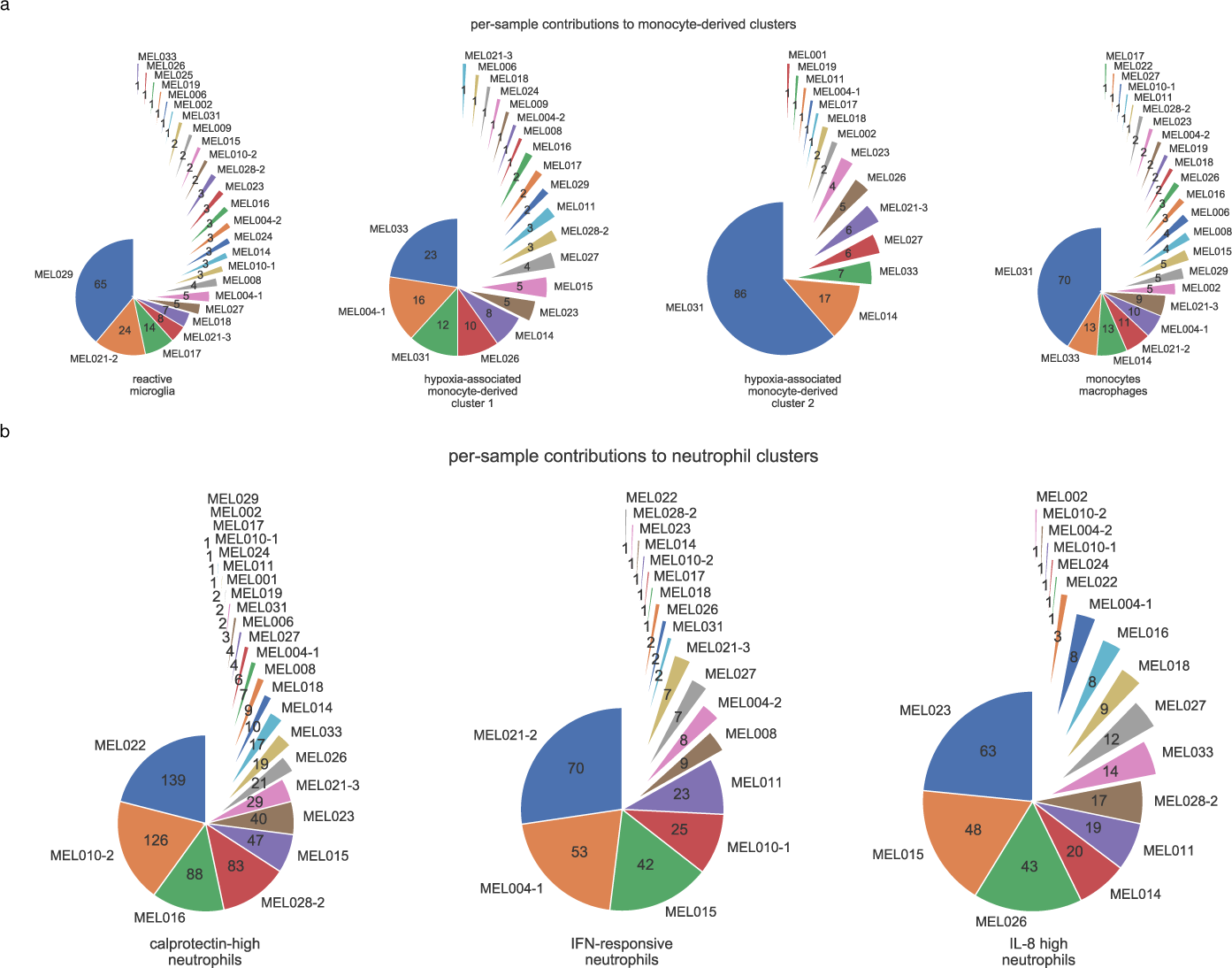
Distribution of myeloid phenotypes across samples. a) Pie chart demonstrating the distribution of macrophage, monocyte, and microglial phenotypes with the absolute number of cells annotated on each slice. b) Pie chart demonstrating the distribution of non-degranulating neutrophil phenotypes with the absolute number of cells annotated on each slice.

**Supplementary Figure 5.**
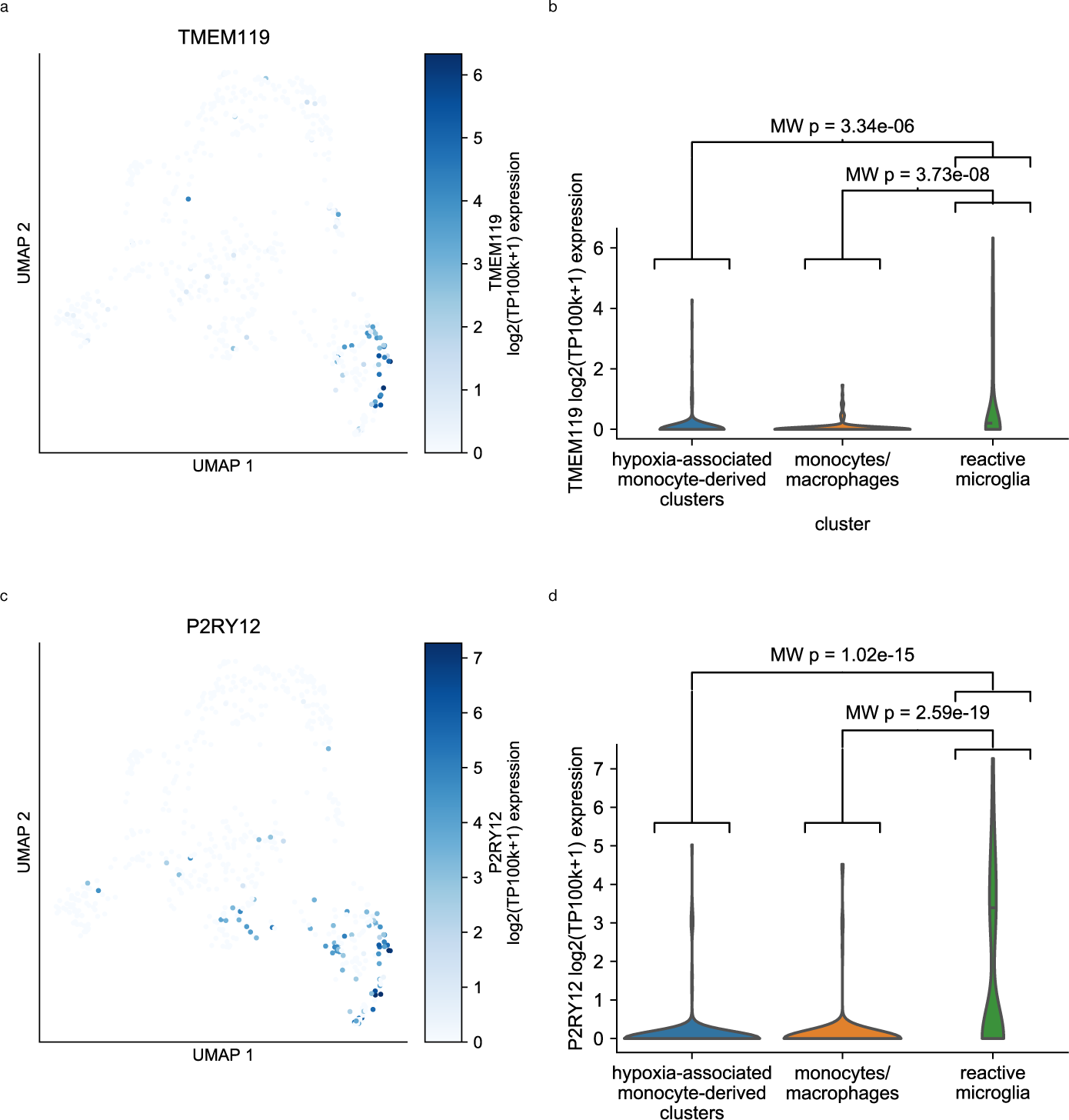
Microglial markers in monocyte-derived cells. a) Levels of microglial marker TMEM119 observed in monocyte-derived cells (UMAP same as that in Fig 2c). Log2(TP100k+1) expression projected as color (colorbar indicated) onto UMAP scatter plot. b) Violin plot showing relative expression (log2(TP100k+1)) of TMEM119 across hypoxia-associated (both hypoxia-associated clusters combined), monocytes/macrophages, and reactive microglia. Two-sided Mann-Whitney U p-values between reactive microglia and other groups indicated. c) Levels of microglial marker P2RY12 observed in monocyte-derived cells (UMAP same as that in Fig 2c). Log2(TP100k+1) expression projected as color (colorbar indicated) onto UMAP scatter plot. d) Violin plot showing relative expression (log2(TP100k+1)) of P2RY12 across hypoxia-associated (both hypoxia-associated clusters combined), monocytes/macrophages, and reactive microglia. Two-sided Mann-Whitney U p-values between reactive microglia and other groups indicated.

**Supplementary Figure 6.**
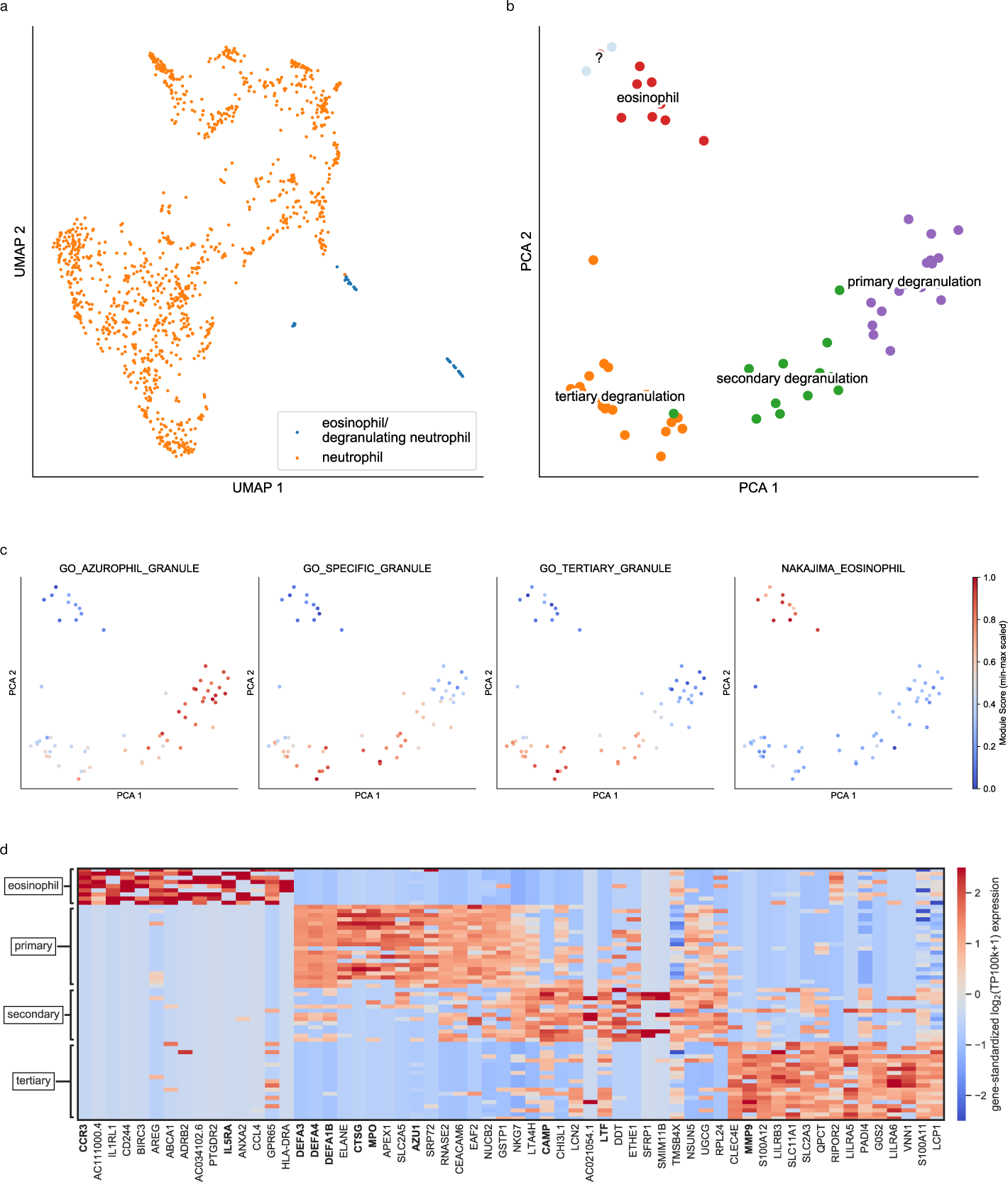
Eosinophil/degranulating neutrophil phenotypes. a) UMAP of neutrophils with eosinophil/degranulating neutrophils, indicating separate clustering of former and latter. b) Distribution in first two principal components of degranulating neutrophils and eosinophils, with clusters annotated. c) UMAP with color map indicating eosinophil and degranulation module scores. d) Heatmap of top 15 marker genes for eosinophil, primary, secondary, and tertiary degranulating clusters, with key genes indicated in bold.

**Supplementary Figure 7.**
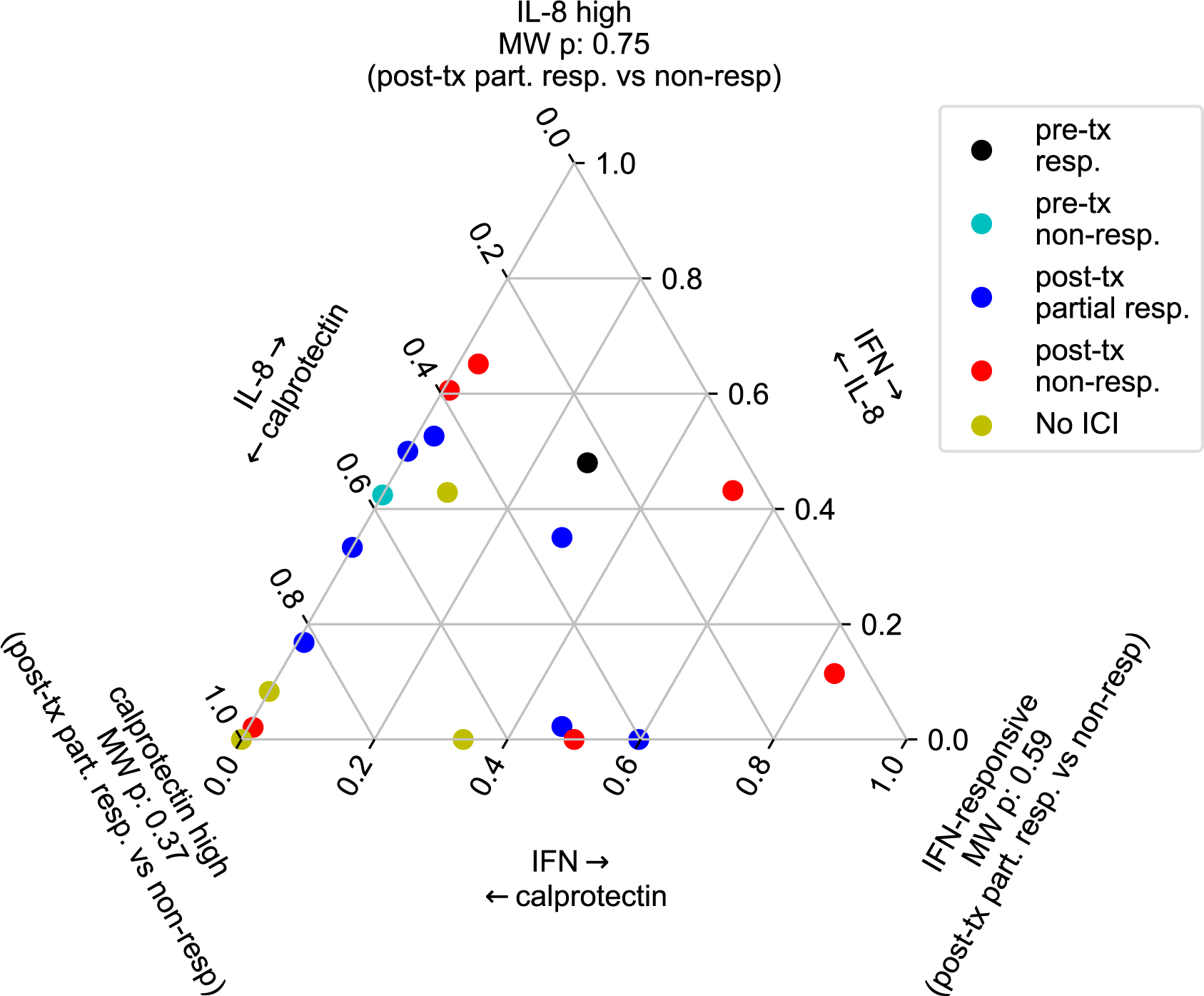
Relative distribution of neutrophil phenotypes. Ternary plot for relative proportion of calprotectin high, IFN-responsive, and IL-8 high neutrophils across patients (values for patients contributing multiple samples are averaged). Mann-Whitney U p-values for the fraction of each population (of all neutrophils) in post-treatment partial vs. non-responder indicated at ternary plot corners.

**Supplementary Figure 8.**
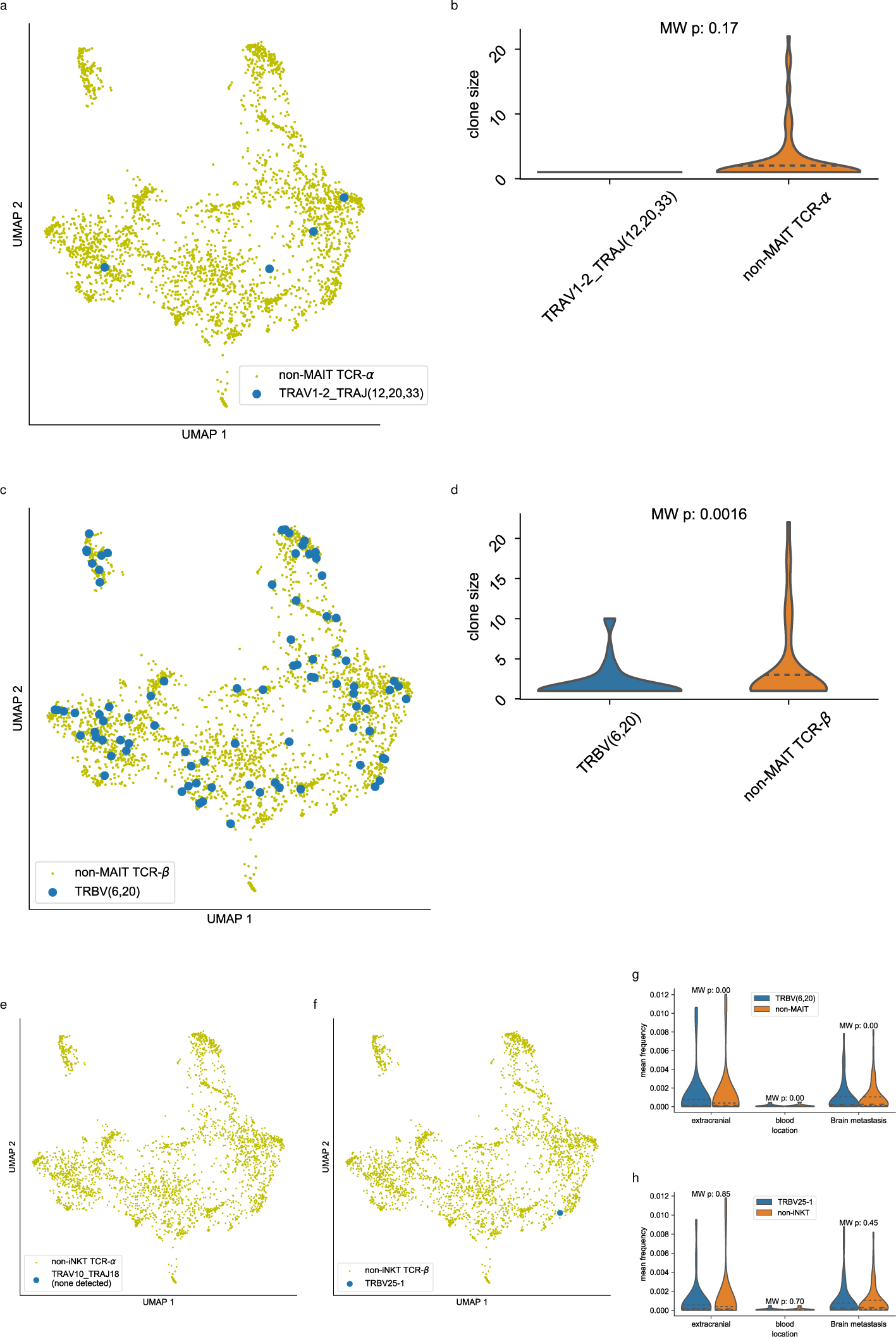
Putative MAIT and iNKT cells. a) UMAP of CD3+ T cells as used in Fig 3a, Fig 4(a,d), with putative MAIT cells (based on presence of TCF-α V allele *TRAV1-2* and TCR-α J allele TRAJ12, TRAJ20, or TRAJ33) indicated. b) Violin plot of clone size of TCR-α based putative MAIT cells vs cells with TCR-α not associated with MAITs. Mann-Whitney U p-value indicated. c) UMAP of CD3+ T cells as used in Fig 3a, Fig 4(a,d), with putative MAIT cells (based on presence of TCF-β V allele *TRBV6* or *TRBV20)* indicated. d) Violin plot of clone size of TCR-β based putative MAIT cells vs cells with TCR-β not associated with MAITs. Mann-Whitney U p-value indicated. e) UMAP of CD3+ T cells as used in Fig 3a, Fig 4(a,d), with putative iNKT cells (based on presence of TCF-α V allele *TRAV10* and TCR- J α allele TRAJ18) indicated. f) UMAP of CD3+ T cells as used in Fig 3a, Fig 4(a,d), with putative iNKT cells (based on presence of TCF-β V allele *TRBV25-1)* indicated. g) Violin plot of mean frequency (within sample) of clonal rearrangements which are putative MAIT cells (based on presence of TCF-β V allele *TRBV6* or *TRBV20)* vs those of clonal rearrangements which are not putatively MAIT cells, separated by tissue type (extracranial, blood, brain metastasis). Mann-Whitney U p-values indicated. h) Violin plot of mean frequency (within sample) of clonal rearrangements which are putative iNKT cells (based on presence of TCF-β V allele *TRBV25-1)* vs those of clonal rearrangements which are not putatively iNKT cells, separated by tissue type (extracranial, blood, brain metastasis). Mann-Whitney U p-values indicated.

**Supplementary Figure 9.**
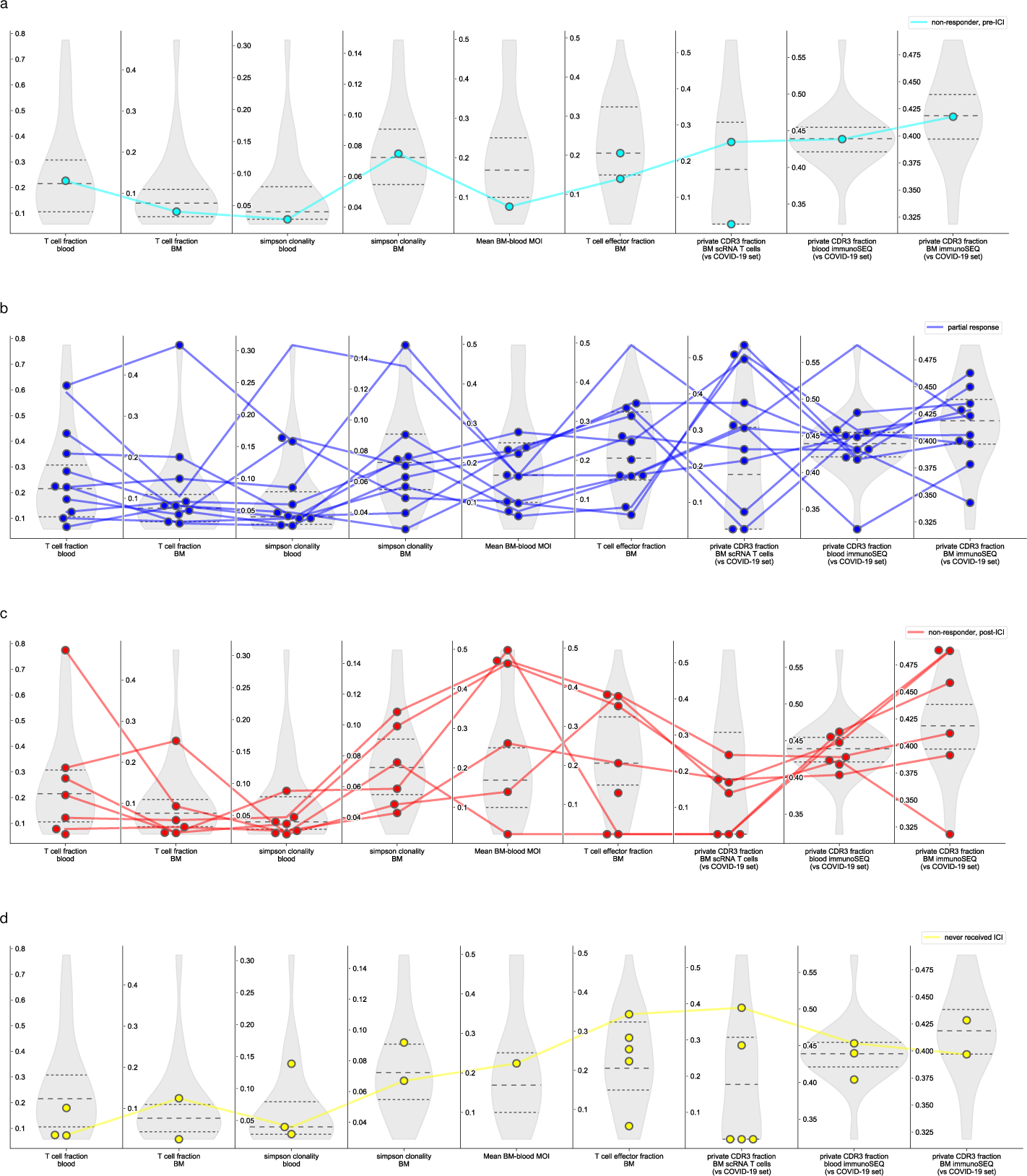
Multiple prognostic metrics across patients. Distribution for each metric (see **Methods**), with upper, lower quartiles indicated by dotted lines, median by dashed line. Line connects values for the same patient for a) pre-ICI individuals who later do not respond to ICI, b) post-ICI individuals demonstrating partial response to ICI c) post-ICI individuals demonstrating non-response to ICI, and d) individuals never receiving ICI. Where individuals are missing values for one or more prognostic metrics, no lines were drawn.

